# Identifying gene regulation modules associated with tumor metastasis using a network decomposition approach and combinatorial fusion analysis

**DOI:** 10.1101/2025.11.17.688950

**Authors:** Aninda Astuti, Christina Schweikert, D. Frank Hsu, Ka-Lok Ng

## Abstract

The hypothesis that modular decomposition of molecular networks into gene regulatory modules (GRMs) enables identification of metastasis-associated gene candidates was systematically evaluated. We developed an efficient bioinformatics approach to identify metastasis-associated GRMs in cancer networks. Using a subgraph method to extract GRMs, we applied combinatorial fusion analysis (CFA) to prioritize them based on relevance. To validate the top-ranked GRMs, we employed the hallmark of cancer annotations, enrichment analysis, drug-target gene evaluation, and survival analysis. The cooperativity effect of GRMs was examined through comparative analysis with previously published studies. Results demonstrate that the proposed approach could effectively identifies metastasis-associated GRMs relative to existing methods.

Robustness was assessed through ten feature combinations and comparisons of three-node versus four-node GRMs. Results consistently confirmed the method’s reliability across different scenarios. This integrated approach combines the subgraph method and CFA to uncover metastasis-associated GRMs effectively. Validation through enrichment analysis, drug-target gene insights, and survival data demonstrates its potential for identifying metastasis-associated target genes and discovering therapeutic drug candidates. Compared to three KIRC cohorts metastasis studies, our approach more effectively identifies GRMs associated with metastasis.

---

Gene cooperativity, where interactions between genes lead to outcomes that are not predictable by single gene effects, plays a crucial role in understanding cancer formation and tumor metastasis. Gene regulatory modules (GRMs) have been constructed to investigate gene interactions, including a lung cancer gene network [1], and tools like FDRnet, which employs Bayesian analysis to identify seed genes and form localized networks [2]. Huang et al. [3] developed a gene target network for lung cancer using the STRING database [4] to map interactions among differentially expressed genes and extract key modules. Gene synergy plays an essential role in tumor metastasis [5], with Xu et al. [6] showing that integrating gene regulation data into graph convolutional networks can effectively construct pan-cancer GRMs, using databases such as HumanNet v2 [7], KEGG [8], BioCarta [9] and Reactome [10] databases.

Interconnected gene modules are capable of performing specific cellular functions, with dynamic motifs like coupled feedback loops [11] enhancing both robustness and evolutionary potential. For instance, positive and negative feedback loops can induce oscillatory cell cycle behaviors [12], while studies on *E. coli* [13] have revealed how coupled feed-forward loops influence protein expression. These regulatory mechanisms are not only critical for normal cellular processes but also play a role in cancer progression.

Tumor metastasis, a primary cause of cancer-related deaths, is one such process where dysregulated gene networks contribute to malignancy. Identifying molecular biomarkers in these GRMs is crucial for advancing targeted therapies and improving diagnostics. Studies have uncovered various crosstalk between metastasis-associated pathways [14, 15] and biomarkers, including microRNA markers in Kidney Renal Clear Cell Carcinoma (KIRC) [16] and gene markers in hepatocellular carcinoma (HCC) [17].

Current research employs multiple strategies to construct GRMs. First, use of protein-protein interactions (PPIs) and cluster analysis to identify metastasis-associated genes [18]. Second, the use of PPIs to infer GRMs and the application of graph theory to analyze network topology through metrics like degree, betweenness, and centrality. Centrality measures have been used to identify oncogenes [19], global regulators [20], glioblastoma biomarkers [21], and genes linked to drug resistance [22]. Third, *Weighted Gene Co-expression Network Analysis (WGCNA)* [23] was applied to detect clusters of highly correlated genes, yet it fails to reveal the regulatory interactions. For example, *WGCNA* has been utilized to identify diagnostic biomarkers for colorectal cancer [24] and markers associated with immune cell infiltration in kidney disease [25]. Fourth, single-cell data have been used in attempts to predict GRMs, these approaches continue to fall short in capturing regulatory relationships [26]. Fifth, multi-omics approach was proposed to construct GRMs for cancer cohorts but still lacking of regulatory information [27]. Sixth, studies are attempting to predict directed GRMs using single-cell multi-omics datasets by integrating scRNA transcriptomic and methylation data to predict GRMs for liver cancer and bladder based on back-propagation neural network [28]. The primary concern with this method is that several parameter choices lack clear justification—specifically, the formulation of the indicator equation, the selection of genes for constructing the final GRMs, and the measurement of gene importance via gene strength.

The above-mentioned approaches typically suggest that directional regulatory relationships between genes cannot be fully captured with high confidence [29]. Instead of replying on *WGCNA* or multi-omics method, we take an *empirical* approach.

This study proposes decomposing molecular networks into smaller GRMs, as developed in our previous work [30], and then ranking their importance with respective to tumor metastasis. This approach has certain advantages, including: (i) interconnected GRMs are experimentally verified, (ii) it can capture the important GRMs in a cancer network and use them as potential drug-targeted modules, (iii) multiple 3-node or 4-node GRMs could be merged to construct n-node GRM, addressing scalability challenges and identifying larger metastasis-associated GRMs.

We hypothesize that the top-ranked GRMs, composed of genetic elements, represent key candidates for metastasis-associated gene modules. To test this hypothesis, we decompose molecular networks into collections of GRMs consisting of four genes (4-node GRMs). Each gene within these modules is assigned four ranking method annotations. We then apply a data fusion approach to prioritize these 4-node GRMs based on their scores.

## By using combinatorial fusion analysis (CFA) to integrate tumor-associated scoring methods, this approach constructs and ranks GRMs associated with metastasis

To further validate our findings, we conducted additional analyses comprising (i) hallmark-of-cancer annotations, (ii) functional enrichment analysis that provides insights into the molecular mechanisms driving tumor metastasis, (iii) drug–target gene evaluation to uncover potential therapeutic targets, and (iv) examination of the cooperativity effects among GRMs.

The next sections under ‘Methods’ will describe the workflow in detail. Then in the ‘Results and Discussions’, we will explore how to scale up from 4-node GRMs to larger modules. For this study, we selected renal cell carcinoma as the focus [31], identifying a set of 4-node GRMs within the KIRC network, each involving interactions between source and target nodes.

The concept of network motifs, introduced by Alon et al. [32], involves comparing a network to a randomized version of itself to identify sub-networks that occur more frequently than expected. However, this approach has limitations, including assumptions about motif frequencies and independence that may lead to false negatives [33]. To address this, we previously proposed a method that does not rely on the null model, using a network subgraph approach instead [30, 34]. Originally referred to as motifs (renamed as subgraphs [30, 34]), we now use the term GRMs to better capture their function.

The study also applies combinatorial fusion analysis (CFA) [35], [36], [37], [38] to overcome challenges in identifying tumor metastasis modules, particularly the lack of upstream and downstream regulatory information. By integrating diverse tumor-associated features, CFA allows for the ranking and prediction of key regulatory modules driving metastasis.

## Methods

In this study, we are aiming to identify tumor metastasis-associated GRMs. We utilized a subgraph method to extract GRMs, followed by CFA for prioritization. Top-ranked GRMs were validated using cancer annotations, enrichment analysis, drug-target gene evaluation, and survival analysis. Cooperativity effects were examined through comparative analysis with existing studies. The workflow diagram for the study is depicted in Figure 1.

**Figure 1.**
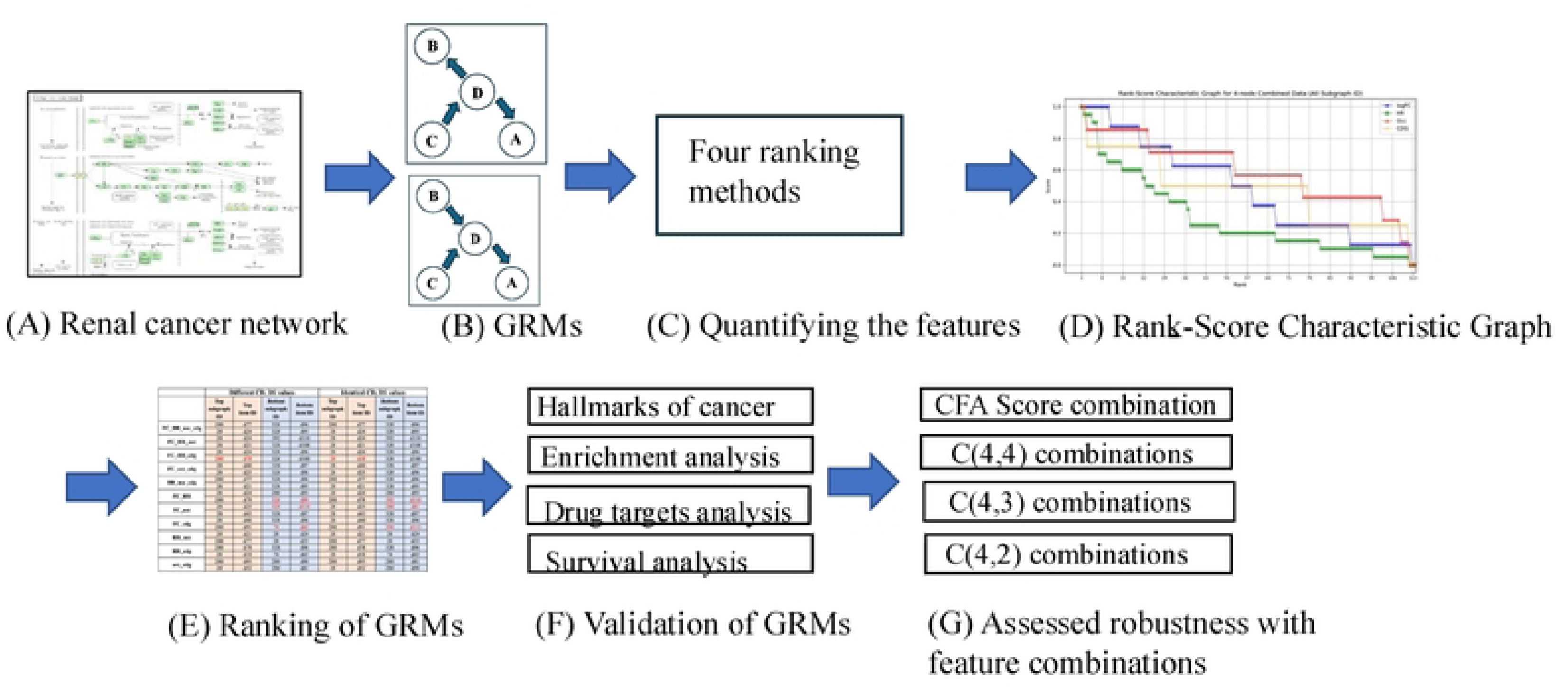
The workflow diagram for the study. (A) Renal cancer network (KIRC) retrieved from the KEGG database. (B) Utilized a subgraph method to extract GRMs. (C) Quantifying the four features. (D) The use of RSC graph for characterizing the four employed scoring methods. (E) Ranking of the GRMs. (F) Validation of GRMs. (G) Accessed robustness with feature combinations.

### Data sources

Molecular biology network data was sourced from the KEGG database, which offers a comprehensive collection of information on biological systems, including genomic, proteomic, chemical, drug, metabolic, and functional data. The cancer cohort used in this study includes M0 and M1 “tumor metastasis” samples from the Xena database (https://xena.ucsc.edu/), which are instrumental in calculating log_2_(Fold Change) (log_2_*FC*) and hazard ratios (*HR*).

To construct and analyze tumor metastasis-associated GRMs, we further evaluated the GRMs using the following additional tumor-associated ranking methods: cancer driver gene (*cdg*) analysis and data from three databases— TMMGdb [39], HCMDB [40], and CMgene [41] — which catalog metastasis-associated genes (*Occ*).

### Utilized a subgraph method to extract GRMs

In previous studies [30, 34, 42], we developed a subgraph approach, an alignment-free method, to analyze 71 molecular networks from KEGG. We decomposed each network into 3-and 4-node GRMs, including the KIRC network, to identify GRMs in the cancer network.

The process followed by the algorithm includes the following steps: (1) First, an adjacency matrix *A*_net_ is constructed for the network to be analyzed, which consists of *N*nodes. The number of nodes incorporated into the subgraph (GRM) is set to four. (2) Four rows and four columns are extracted from *A*_net_, generating 𝐶(*N*, 4) possible combinations. These combinations form a new adjacency matrix, denoted as 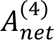 . During this step, it is crucial that the row and column indices align appropriately to form 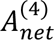, and any disconnected subgraphs are removed. The total number of edges 𝐸within these subgraphs can vary, based on prior research study [43, 44]. For example, a connected 4-node subgraph would have a minimum of three edges, i.e., 𝐸 = 3. (3) The matrix 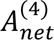 is then converted into integer values, and the number of edges 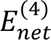 present in this subgraph is counted. (4) This integer value typically represents an isomorphic form of a 4-node subgraph (GRM). The algorithm iteratively applies different permutation matrix operations until it identifies the smallest possible integer, which is then assigned a unique identifier labeled as 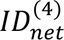 . (5) To identify whether a matching subgraph exists based on its integer representation, subsets of integers corresponding to subgraphs with 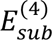 edges (where 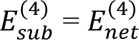) must be selected. The integer identifier 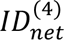 is then compared with the corresponding identifier 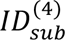 of any potential subgraph 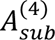. A match between these integers confirms the detection of a 4-node subgraph (GRM) within the original matrix *A*_net_. For a detailed explanation of the method, please refer to the prior research [42].

### Description of the four ranking methods

The first two ranking methods— log_2_*FC* and *HR* score—were derived from our previous work [39]. TMMGdb uses DESeq2 [45] and Cox analysis to calculate log_2_*FC* and *HR* by comparing RNAseq expression data between non-metastatic (M0) and metastatic (M1) samples. There is no universally accepted cutoff for |log₂FC|; thresholds can vary, with some studies using as low as |log₂*FC*| ≥ 0.20 [46] while others apply |log₂*FC*| ≥ 0.58 [47, 48]. In this study, we adopt a threshold which is approximately equals to the average value; |log₂*FC*| = 0.38.

We filtered the data by including any values with |log_2_*FC*| ≥ 0.38 and using an adjusted p-value < 0.05 calculated by DESeq2. The remaining data were then used to establish a threshold that retained 86% of the original dataset. Next, we divide the |log_2_*FC*| range of 0.38 to 0.86 into 10 equal intervals. Scores from 1 to 10 are assigned accordingly, with values greater than or equal to 0.86 receiving a score of 10. The total score of the GRM reflects its significance, with higher scores indicating modules more relevant to tumor metastasis.

The second method ranks GRM genetic elements based on their *HR* score. The *HR* for each gene with *HR* ≥ 1.05 and an adjusted p-value < 0.05 from Cox proportional hazards analysis. Data with *HR* < 1.05 are excluded, and the remaining values are divided into 10 intervals, assigning a score from 1 to 10. If *HR* is less than 1.05, a score of 0 is assigned. Higher *HR* scores suggest a stronger association with tumor metastasis.

The next two methods include documented genetic information. The third method involves verifying whether a genetic element appears in tumor metastasis gene databases. Each gene is checked across three specific databases: TMMGdb, HCMDB, and CMgene, which catalog 4,243, 1,938, and 2,040 metastasis-associated genes, respectively. The module’s score ranges from 0 to 12, with a maximum of 12 indicating that all genes are listed in all three databases.

The fourth method evaluates whether any nodes are cancer driver genes. This analysis determines whether the 4-node module plays a key role in tumor metastasis. The cancer driver gene score ranges from 0 to 4.

For clarity, we use *FC, HR, Occ*, and *cdg* to represent log_2_*FC*, hazard ratio, the frequency of occurrence in the three metastasis gene databases, and cancer driver gene respectively.

### The CFA method in evaluating GRMs

After scoring the GRMs, the next step is to rank them based on the accumulated scores. This process is complicated by the need to combine continuous and categorical values from multiple criteria. To address this, we employ the combinatorial fusion analysis (CFA) framework, which facilitates the combination of different scoring methods. CFA simplifies the complexity arising from diverse scoring criteria and has been applied in various domains such as drug discovery [49], protein structure prediction [50], and the classification of documents according to the United Nations Sustainable Development Goals (SDGs) [51]. It has also been used to enhance data synthesis and prediction quality in the study of metal halide perovskites [52].

Combining multiple scoring systems can be achieved through at least two approaches: score combination (*SC*) or rank combination (*RC*). Let *M* represent a set of *n* scoring methods, *M* = {*A_1_, …, A_i_,…,A_n_*} where *A_i_* denotes the *i*-th scoring method. Consider a dataset *D* comprised of *m* items, *D* = {*d_1_, …, d_k_, …,d_m_*}. Each scoring method A∈M, considered here as a scoring system A, is described by a score function s_A_, rank function r_A_, and a rank-score characteristic (RSC) function f_A_. The score function for a system A, 𝑠*_Ai_*(𝑑_α_), produces a real number or integer. Here, 𝑠*_Ai_*(𝑑_α_), represents the score assigned to item 𝑑_+_by the *i-*th scoring method, A_i_.

Different scoring methods utilize varying metrics; therefore, it is crucial to establish standardized measures and use the normalized score functions for each method. The normalized score function 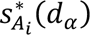 for item 𝑑_α_ using the *i-th* scoring method is defined as:

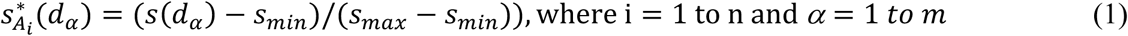

where *𝑠_min_* and *𝑠_max_* represent the minimum and maximum scores, respectively. Normalizing the scores facilitates an effective comparison and integration of the different scoring methods, as they are each now within the range of 0 to 1.

The rank function for scoring system *A*, 𝑟*_A_*, is generated by ordering the normalized score function 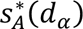 in descending order and mapping the scores to ranks starting at 1 for the highest score. A rank-score characteristic (RSC) function for scoring system *A*, *𝑓_A_*, is a mapping from ranks to scores and is defined as:

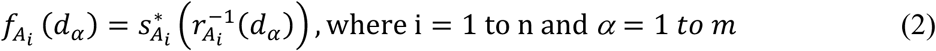

After selecting *n* scoring methods, the average score combination method for data d_α_ is given by:

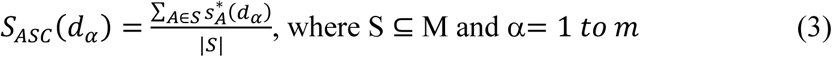

𝑆*_ASC_*(𝑑_α_) indicates that the score for data 𝑑_α_ is the average of the scores from *n* scoring methods.

The aforementioned method involves averaging the scores from four different scoring methods, each with equal weight. However, this approach may introduce bias, as it doesn’t account for the varying degrees of differences among the scoring systems. One solution is to employ cognitive diversity (*CD*) [53, 54] to compute a metric that can be used as a weight for the scoring systems. The *CD* metric uses the area between the rank-score characteristic (RSC) functions of two scoring systems to represent the diversity. The *CD* metric quantifies the difference in scoring behavior between two scoring systems, which is distinct from the Pearson correlation in classical statistics which compares two sets of scores.

The cognitive diversity (*CD*) between two scoring methods, *A* and *B* is computed using the formula in Equation 4.

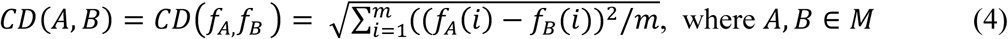

The greater the *CD (diversity)* between two scoring systems, the larger the difference in their scoring behavior. Subsequently, we calculate the diversity strength (*DS*) metric, which represents the average *CD* between the scoring method *A* and other scoring methods, calculated using the following formula:

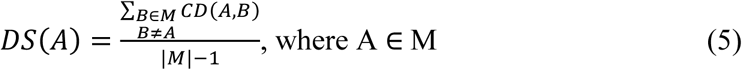

We observed that GRMs frequently share the same ranking for the cancer driver gene (*cdg*) score due to the score’s range of 0 to 4, which often results in identical ‘*cdg*’ scores for different GRMs. Consequently, we have decided to focus on the weighted score combination method. The weighted score combination 𝑆_𝖶𝑆𝐶_(𝑑_𝛼_) can be obtained from the score function 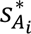 using *DS* as the weight, which is defined by Equation 6 below

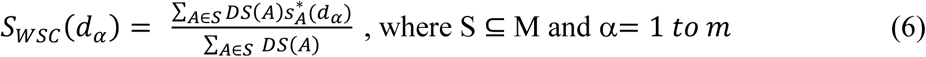

For this study, we will employ the aforementioned methods to rank the GRMs. In summary, combinatorial fusion analysis (CFA) is a framework that uses the Rank-Score Characteristic (RSC) function to measure *CD* between two scoring systems and provides methods for scoring system combinations. *CD* and *DS* measure the dissimilarity or inconsistency between scoring methods.

### Applying multiple score combinations to rank the GRMs

An important step in the analysis is combining the four existing scoring methods. This approach has been explored in previous studies to improve multivariate or multi-objective classification, prediction, and optimization [54–56]. There are 11 possible combinations of the four ranking methods, each yielding a weighted score combination (*SC*) by *DS*. Using KIRC as an example, we construct a 114 x 11 matrix where each row corresponds to a GRM, and each column represents the rankings derived from one of the 11 combinations. A second matrix is created where rows indicate rankings from 1 to 114, and columns show rankings from the 11 models. Modules that rank within the top five across these combinations are selected for further analysis.

### Assess the relevance of the GRMs related to KIRC using GSEA and GOEA

To validate the significance of the ranked 4-node GRMs, we retrieve direct protein-protein interaction (PPI) partners for each genetic element from the STRING database. Enrichment analysis is then performed using DAVID, focusing on the Wikipathwayss and DisGeNET annotations. We examine 4-node GRMs using rankings from both average and weighted score combinations, excluding rank combinations where some methods only allow a limited number of ranks, such as the cancer driver gene ranking.

The Molecular Signatures Database (MSigDB) [57] organizes gene sets into collections, including the cancer-related hallmark gene set. For this study, we used the hallmark gene set from MSigDB (v2024.1.Hs, August 2024), which includes 50 gene sets. Gene Set Enrichment Analysis (GSEA) is employed to analyze the significance of the GRMs in relation to these hallmark pathways.

The Wikipathwayss annotation in DAVID provides access to biological pathways curated by the scientific community, helping to visualize and analyze gene functions and interactions within these pathways. Additionally, the DisGeNET database [58] supports the identification of gene-disease associations, offering valuable insights into the genetic basis of human diseases from a variety of sources, including expert-curated repositories and GWAS catalogs.

### Assess the relevance of the GRMs related to KIRC using drug-target information and survival analysis

Determining whether GRMs are involved in critical signaling pathways is essential, as their dysregulation can lead to uncontrolled cell growth and cancer formation. The KEGG drug database (https://www.genome.jp/kegg/drug/) is used to assess whether the GRMs are associated with known drug compounds. This resource includes information on approved drugs in Japan, the USA, and Europe, and links drugs to their therapeutic targets, molecular interactions, and drug metabolism, which can serve as potential biomarkers for diagnosis and prognosis.

The survival analysis results for the top and bottom ranked GRMs are shown in the Kaplan-Meier (KM) plots, which are derived from queries in the UALCAN database [59, 60].

## Results and Discussions

### The Rank-Score Characteristic (RSC) Graph, cognitive diversity and diversity strength

Upon decomposing the KIRC network into 4-node GRMs, a limited variety of GRM types were identified; that is, GRMs with large graph energies are not present [30, 34], rather than the full spectrum of 199 possibilities. The specific GRMs found include types: {d_1_, d_2_, …, d_11_} = {14, 28, 74, 76, 92, 280, 328, 344, 392, 394, 2184}, as detailed in Supplementary File 1. The complete topology of all potential 4-node GRMs is available in Supplementary File 2.

Should any genetic element within a GRM be without a feature score value, the GRM is excluded, leaving 114 for analysis. Our previous research [30] noted the absence of certain GRM patterns, with the missing types linked to significant graph energies. Networks of molecular interactions are built from a finite number of regulatory topologies.

We opted for a thorough examination of the 4-node GRMs over the 3-node counterparts due to the nearly doubled quantity, offering more detailed insights. Figure 2 presents the RSC graph for the four employed scoring methods.

**Figure 2.**
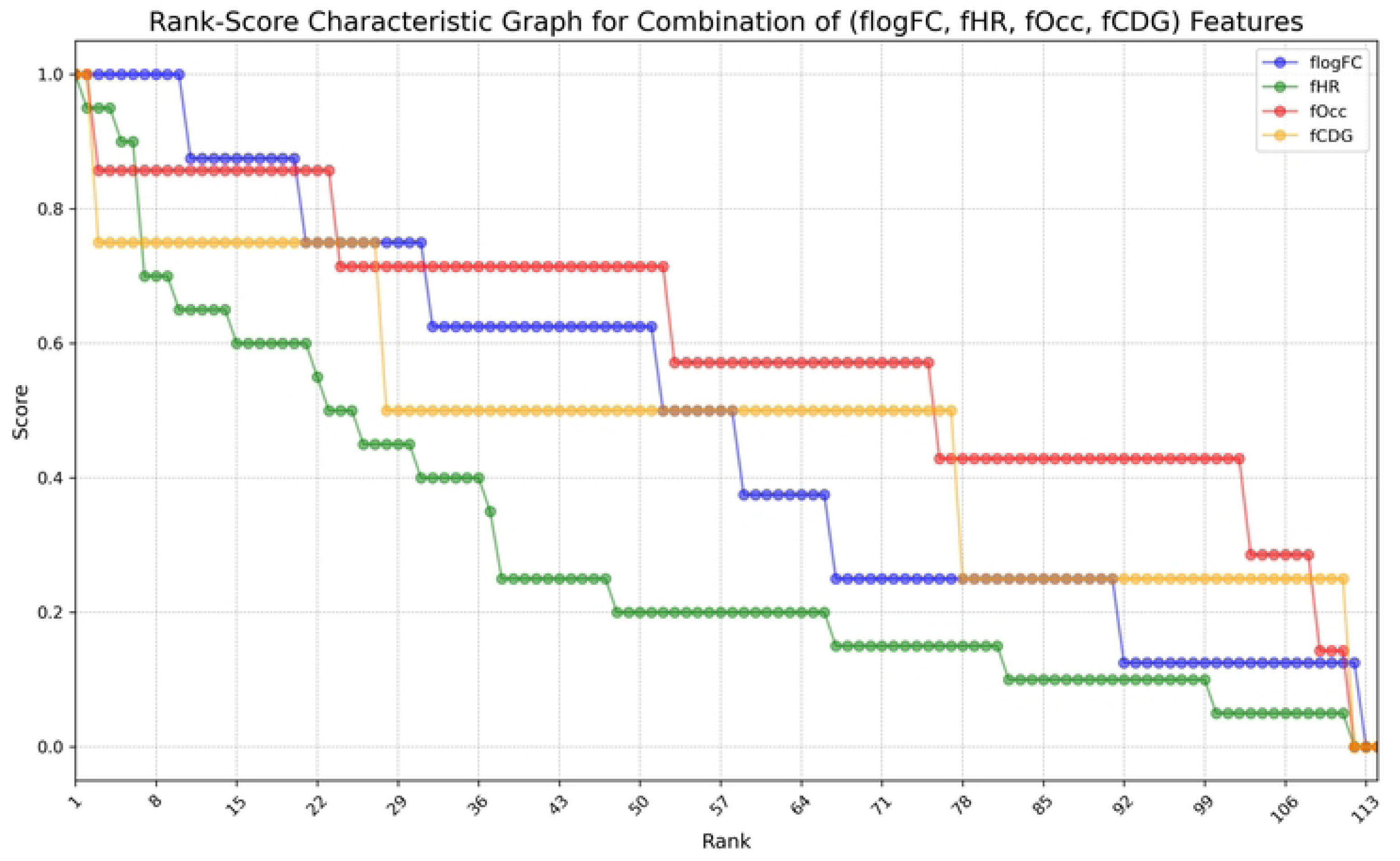
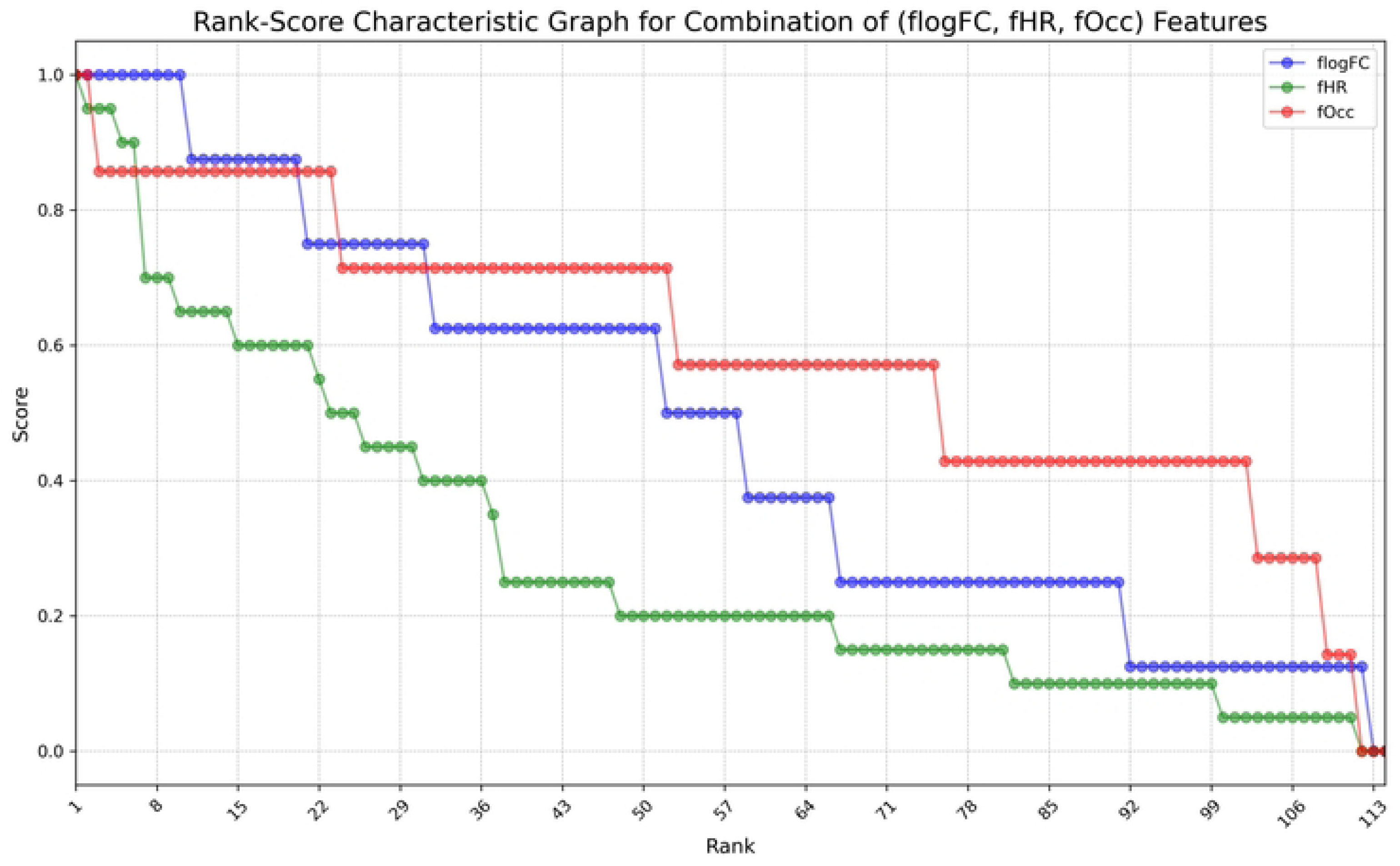
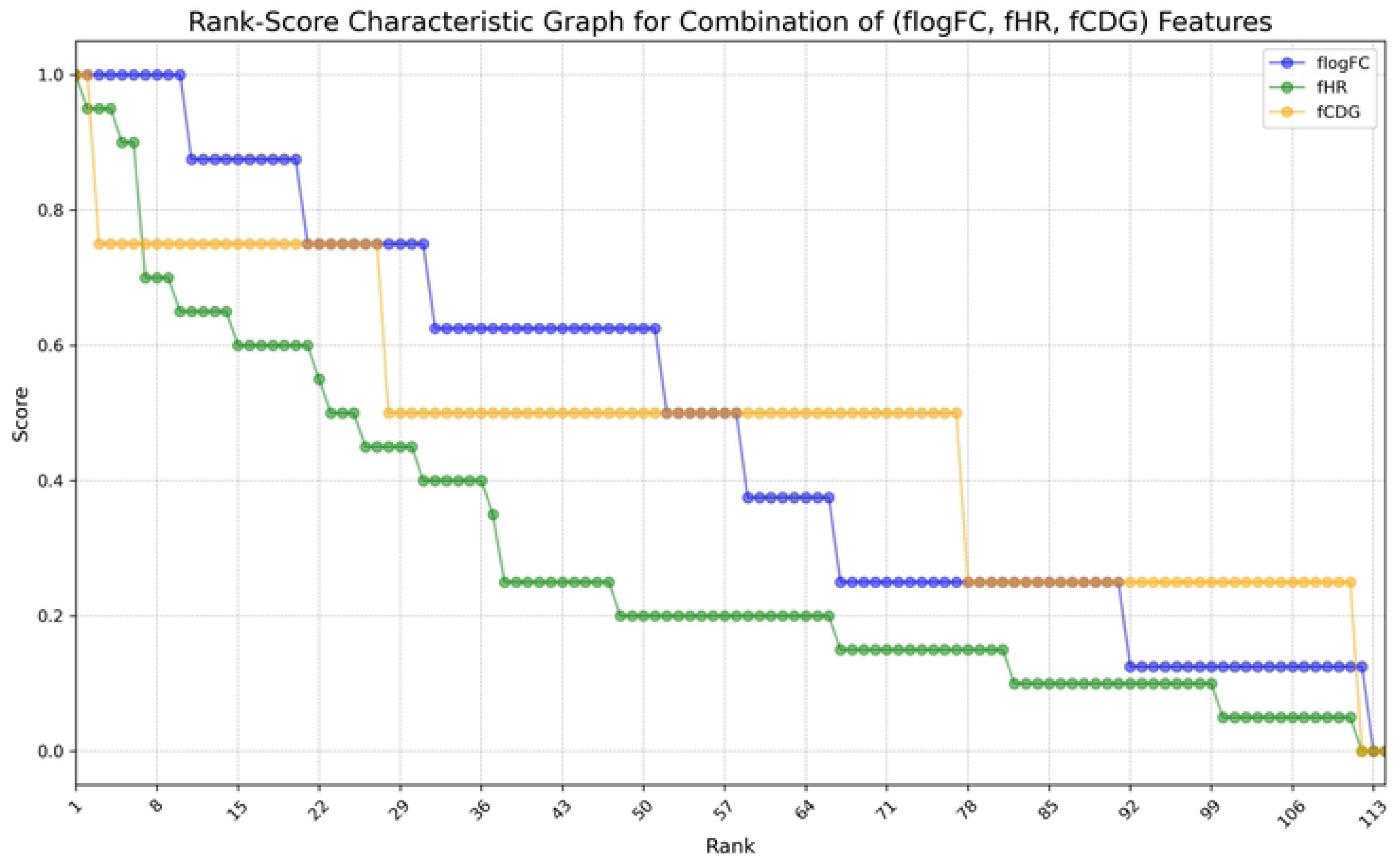
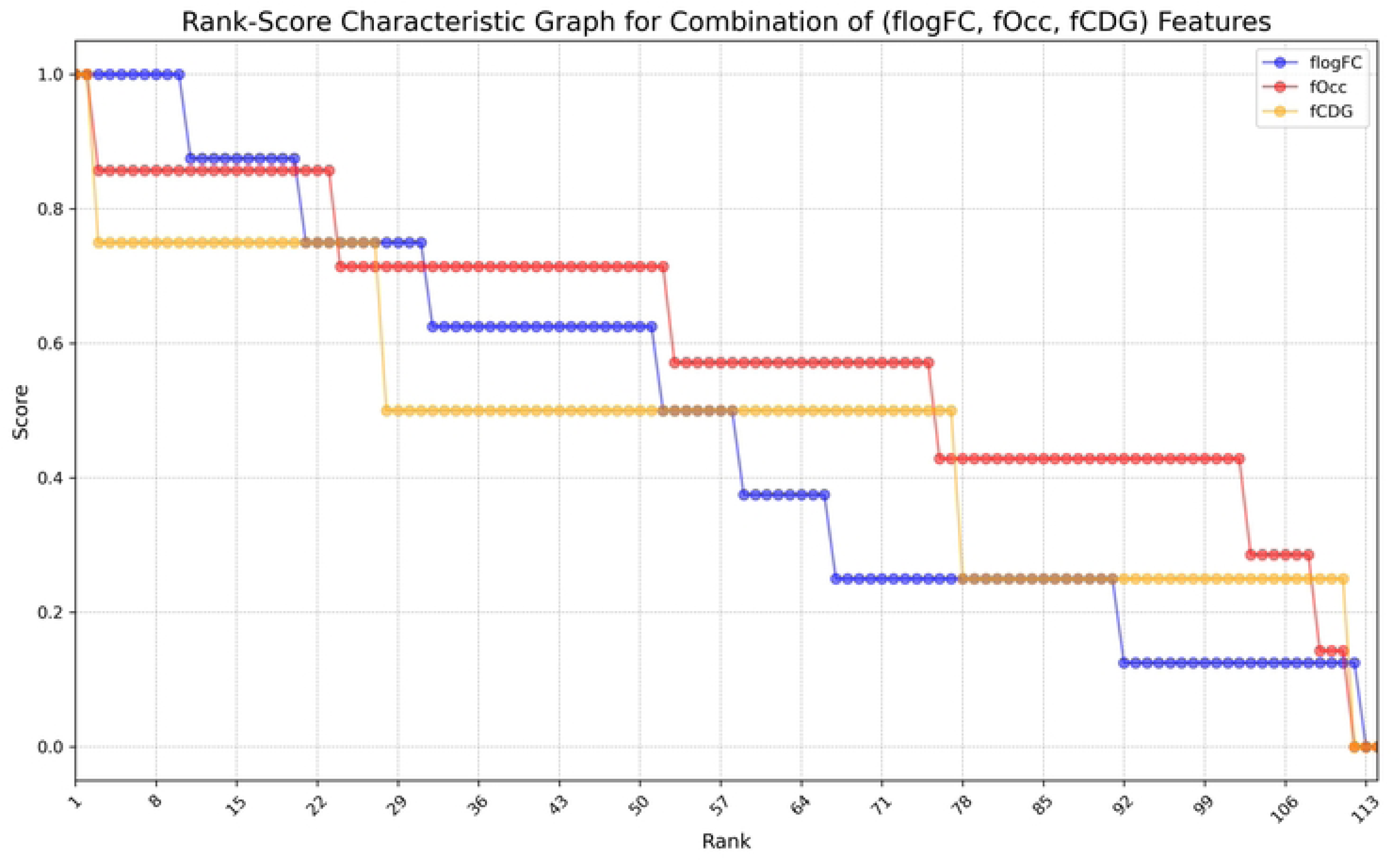
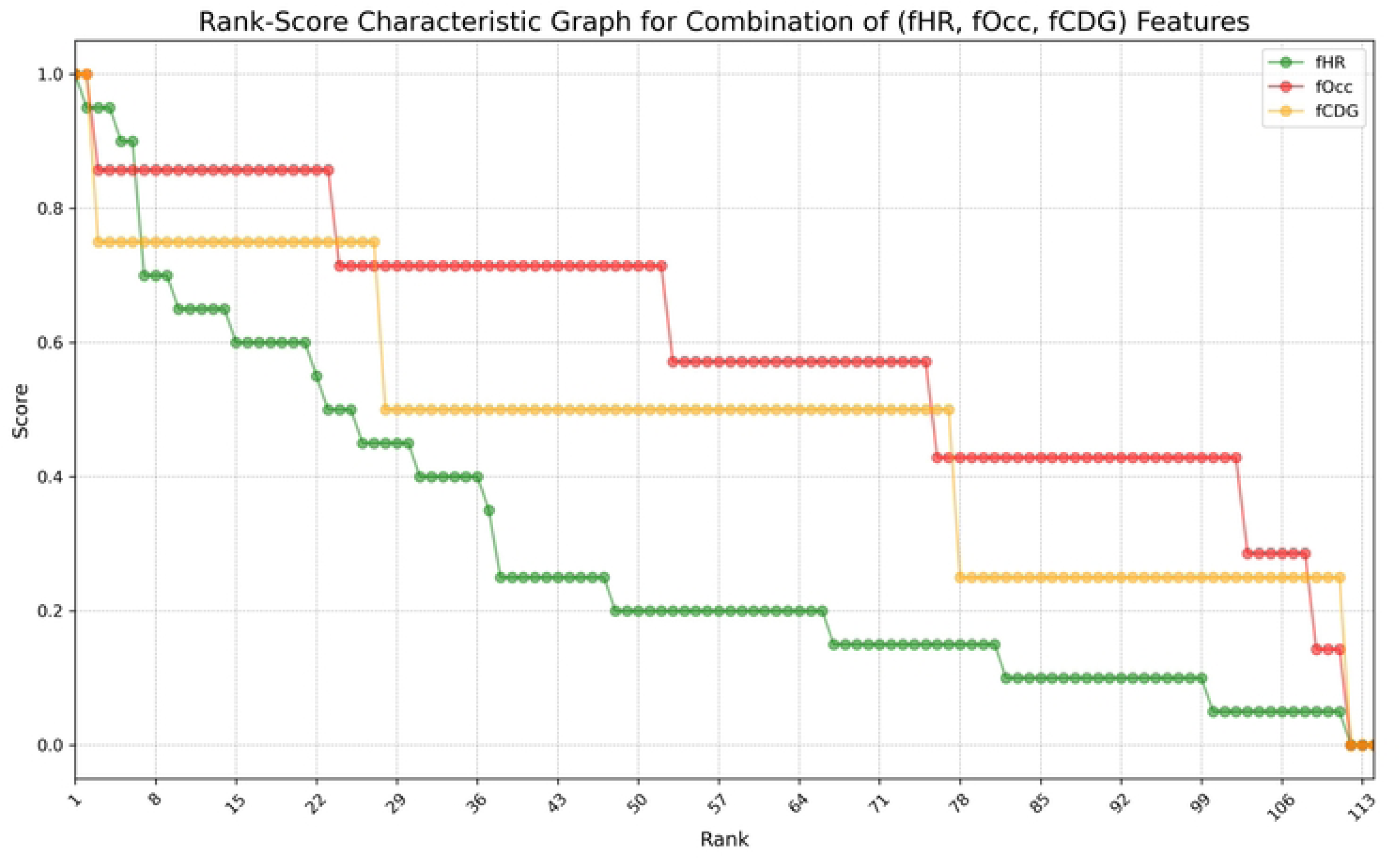
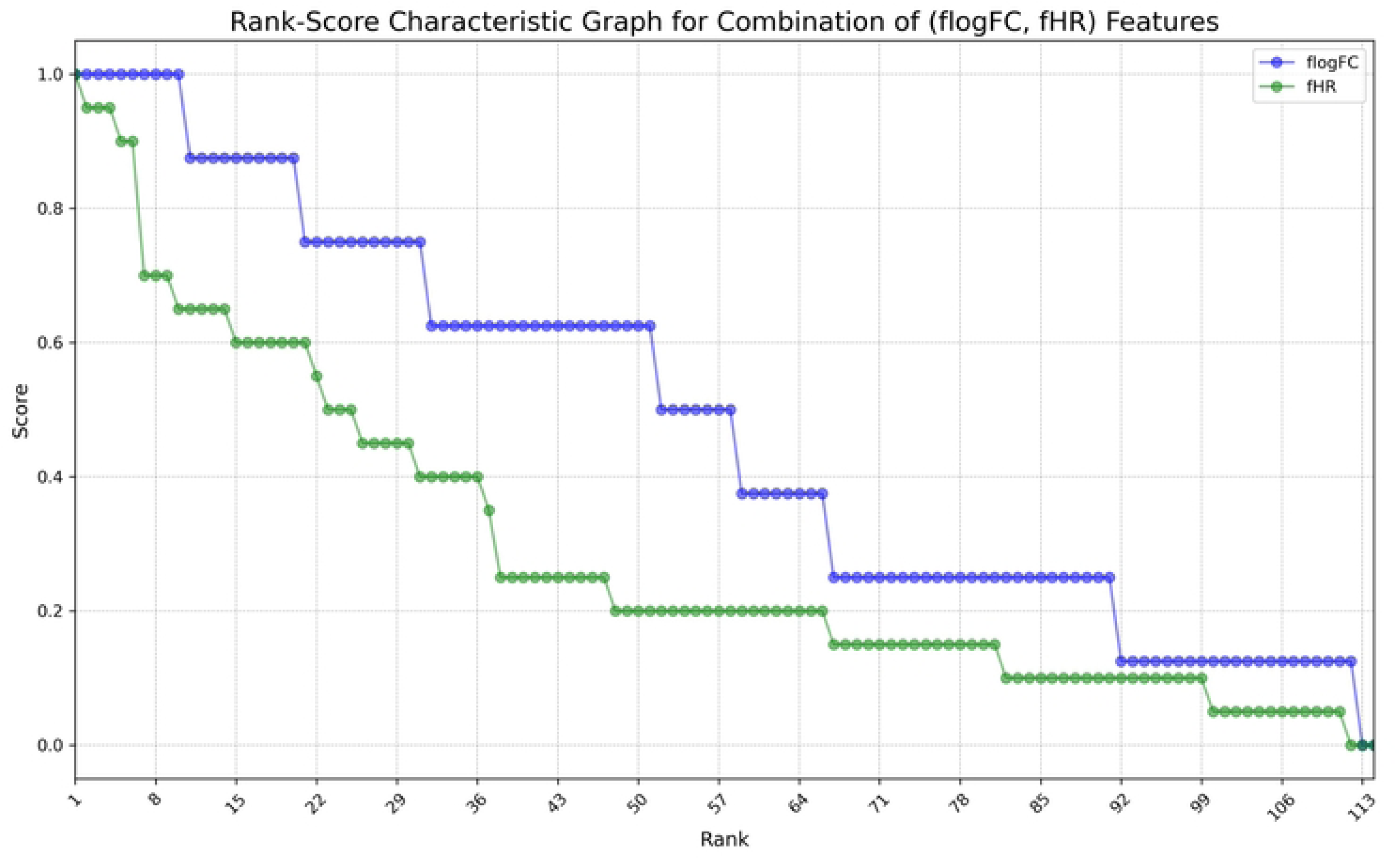
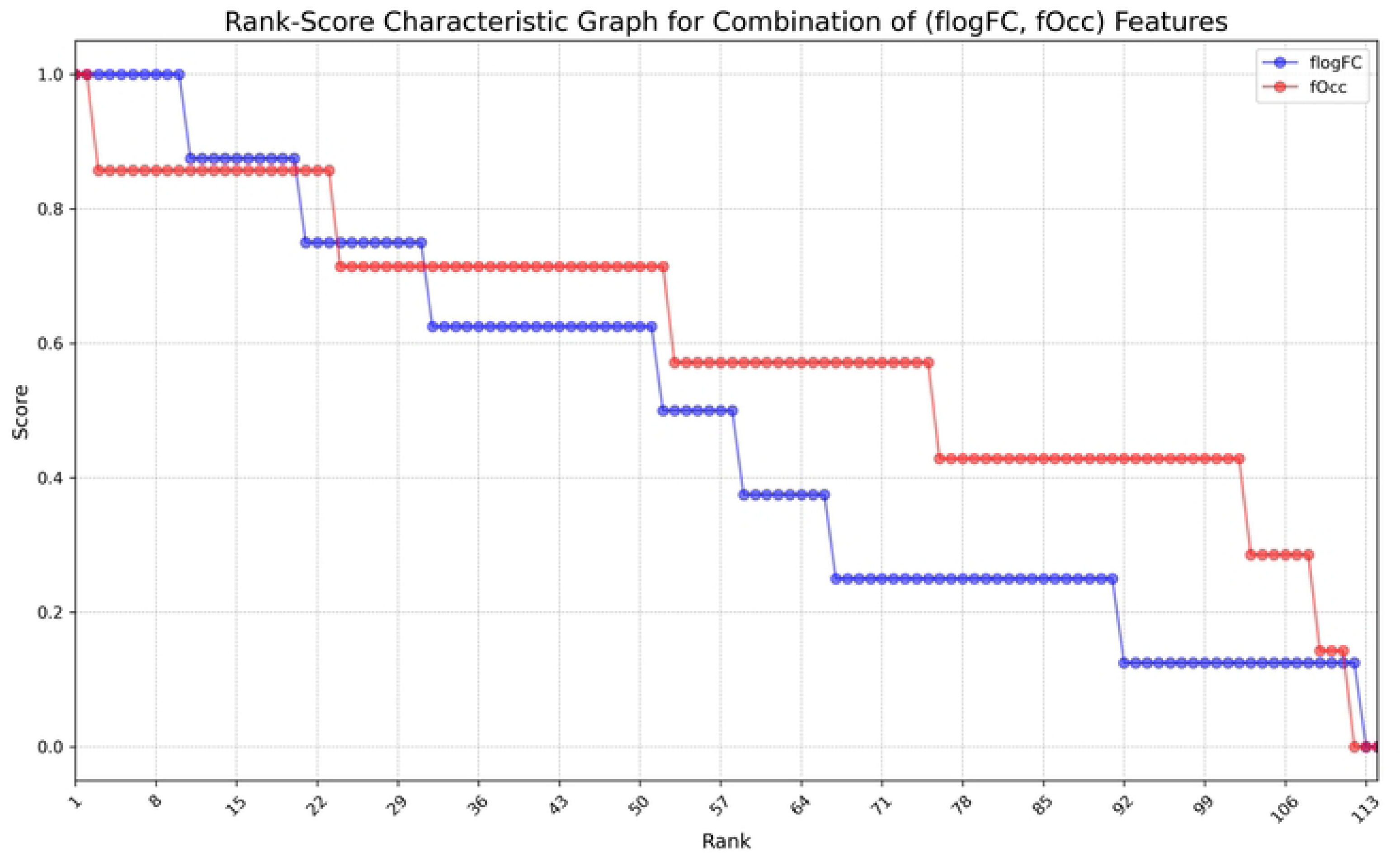
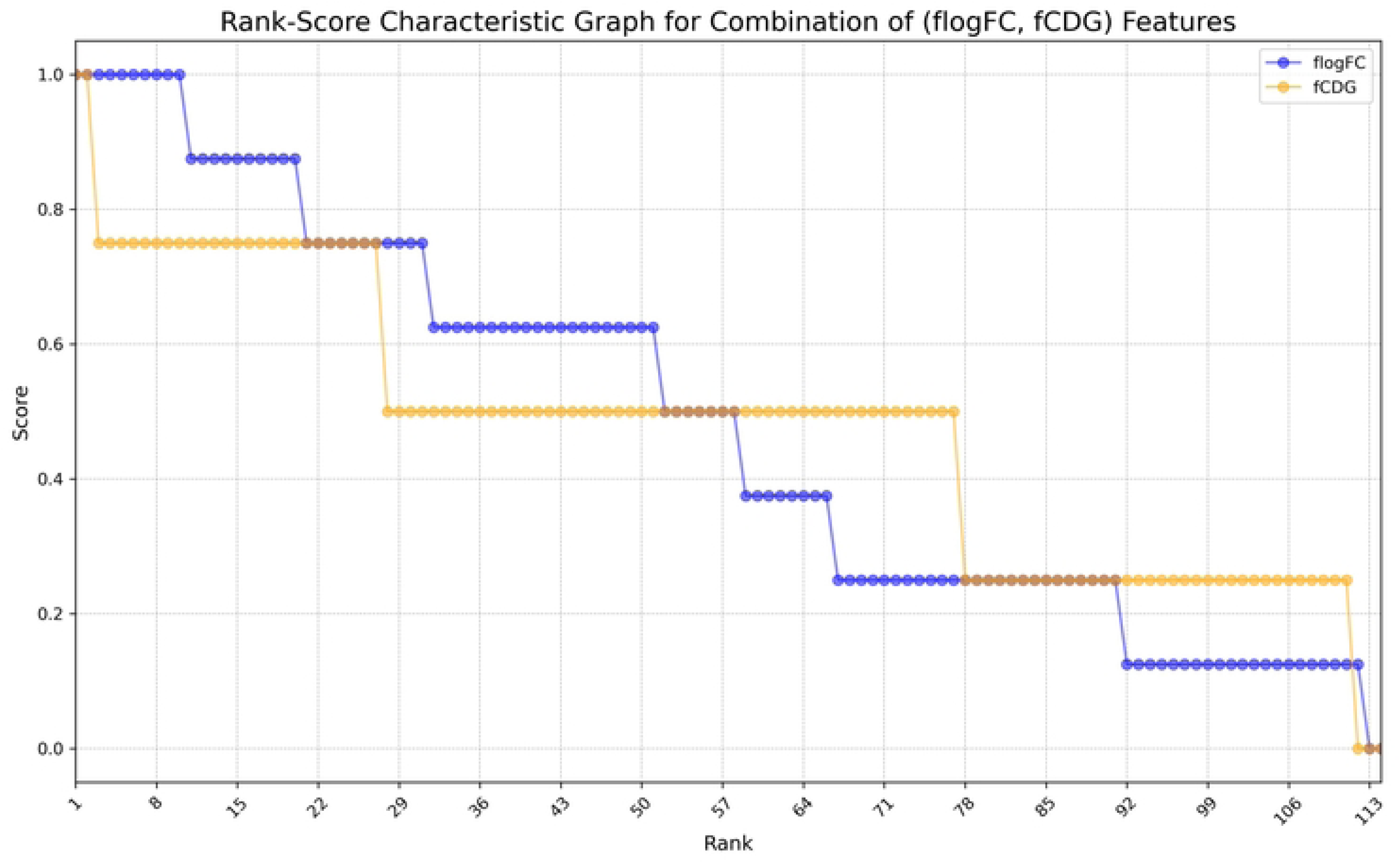
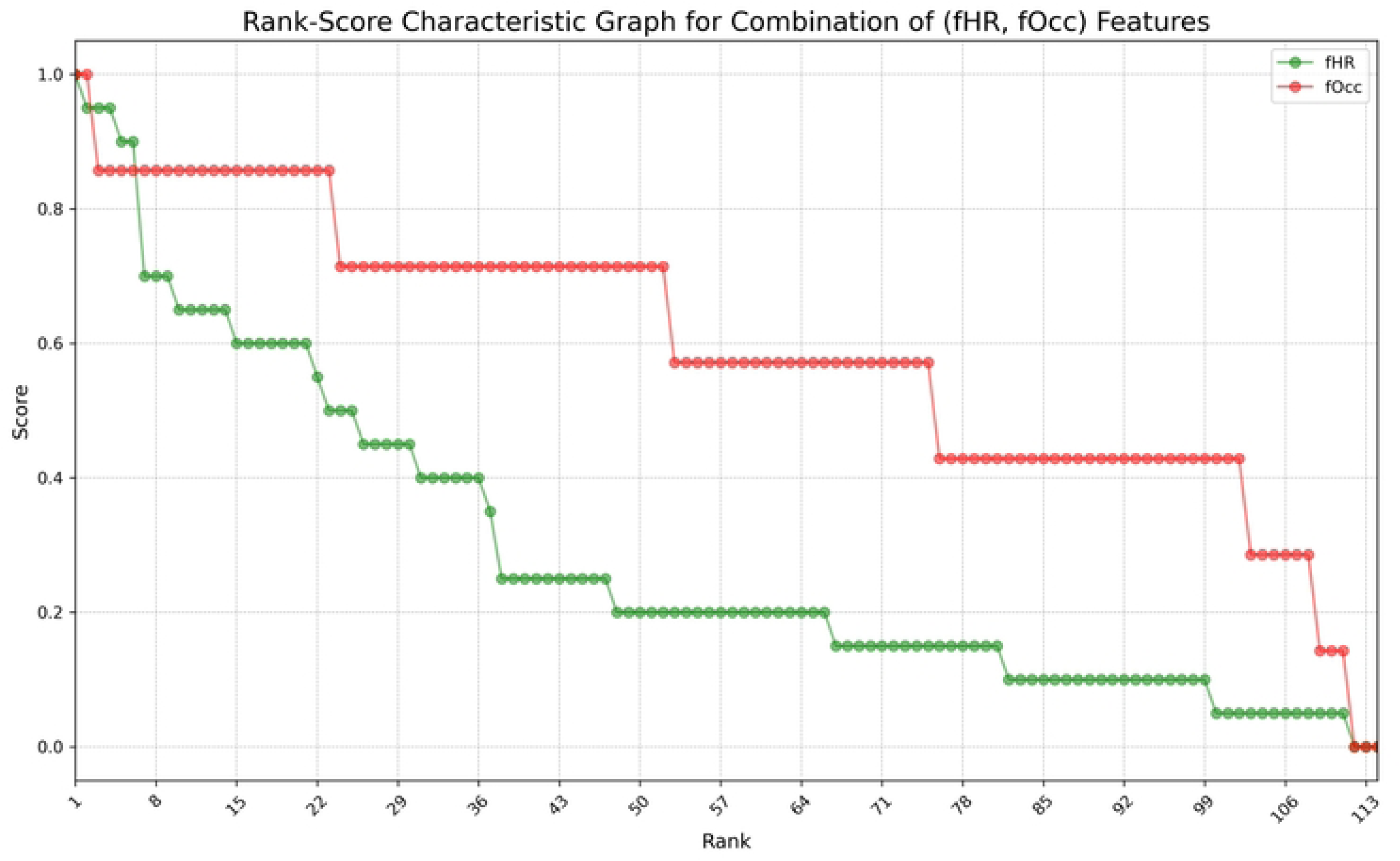
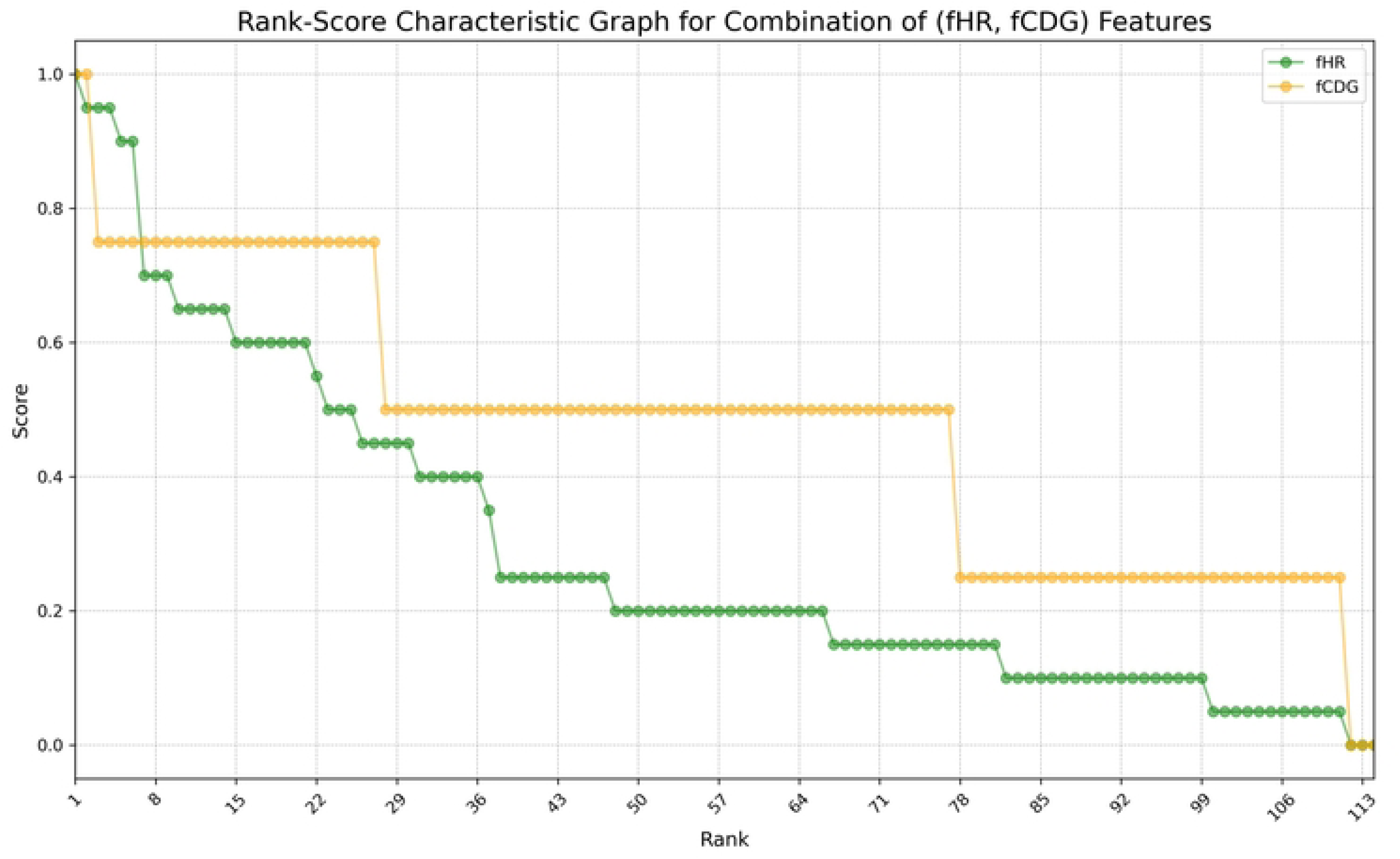
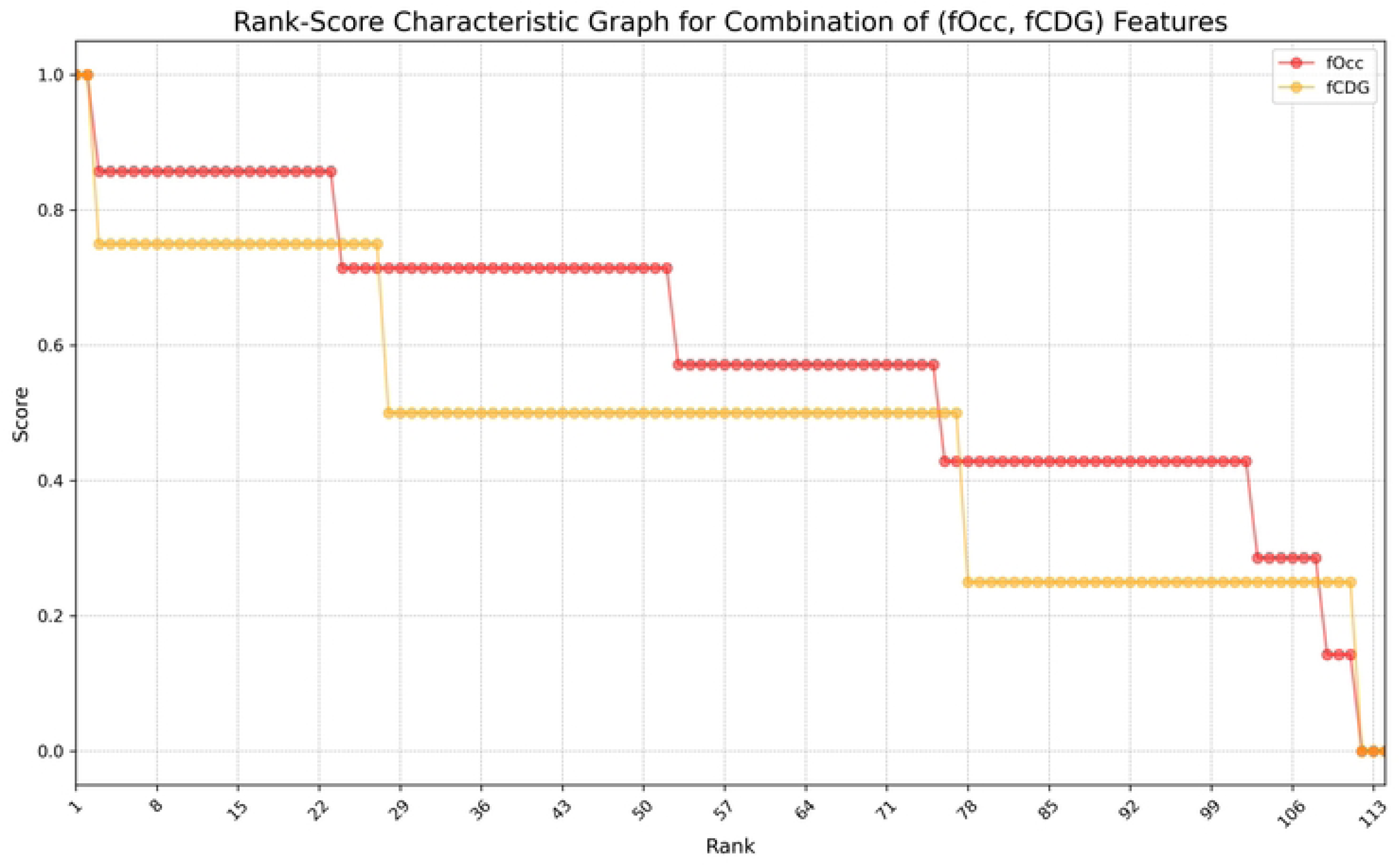
The Rank-Score Characteristic Graph for the 11 combinations of scoring methods.(A) The four features *log(FC), HR*, *Occ* and *cdg*. (B) *log(FC), HR* and *Occ*. (C) *log(FC), HR* and *cdg*. (D) *log(FC)*, *Occ* and *cdg.* (E) *HR*, *Occ* and *cdg.* (F) *log(FC)* and *HR*. (G) *log(FC)* and *Occ*. (H) *log(FC)* and *cdg.* (I) *HR* and *Occ*. (J) *HR* and *cdg.* (K) *Occ* and *cdg*.

Table 1 outlines the *CD* scores derived from the six possible combinations of two out of the four scoring systems. The greatest distance observed is 0.329 between the *HR* and *Occ* plots, suggesting these methods assess GRMs differently. In contrast, the smallest distances are both 0.144, found between the *cdg* and *FC* plots and the *cdg* and *Occ* plots, indicating similar assessments by these methods.

**Table 1.** The *CD* values for the six possible pairs of models, where *CD(X,Y)* represents the diversity between ranking methods *X* and *Y*, where *X* and *Y* are elements of {*FC* (equivalent to *log(FC)), HR, Occ*, *cdg*}.

| $CD(X,Y)$ | $CD(FC,HR)$ | $CD(FC,Occ)$ | $CD(FC,cdg)$ | $CD(HR,Occ)$ | $CD(HR,cdg)$ | $CD(Occ,cdg)$ |
| --- | --- | --- | --- | --- | --- | --- |
| $CD$ value | 0.223 | 0.172 | 0.144 | 0.329 | 0.219 | 0.144 |

We then computed the *DS* metric to represent the average *CD* across different scoring methods. Supplementary Table 1 lists the *DS* values for 11 possible scoring methods, with *DS_Z* denoting the DS between scoring method Z and the other scoring methods.

The *HR* feature’s *DS* value is the largest at 0.257, suggesting its distinct rating approach. This is supported by the *DS_HR* column in Supplementary Table 1, which shows the largest distances. When comparing only two scoring methods, the *DS* value is symmetric: *DS(X, Y) = DS(Y, X)*.

Table 2 compares weighted score combination (*SC*) by *DS* and average score combination in GRM ranking, focusing the top two and bottom two ranked GRMs, the rationale being to uncover GRMs exhibit different biological functions.

**Table 2.**
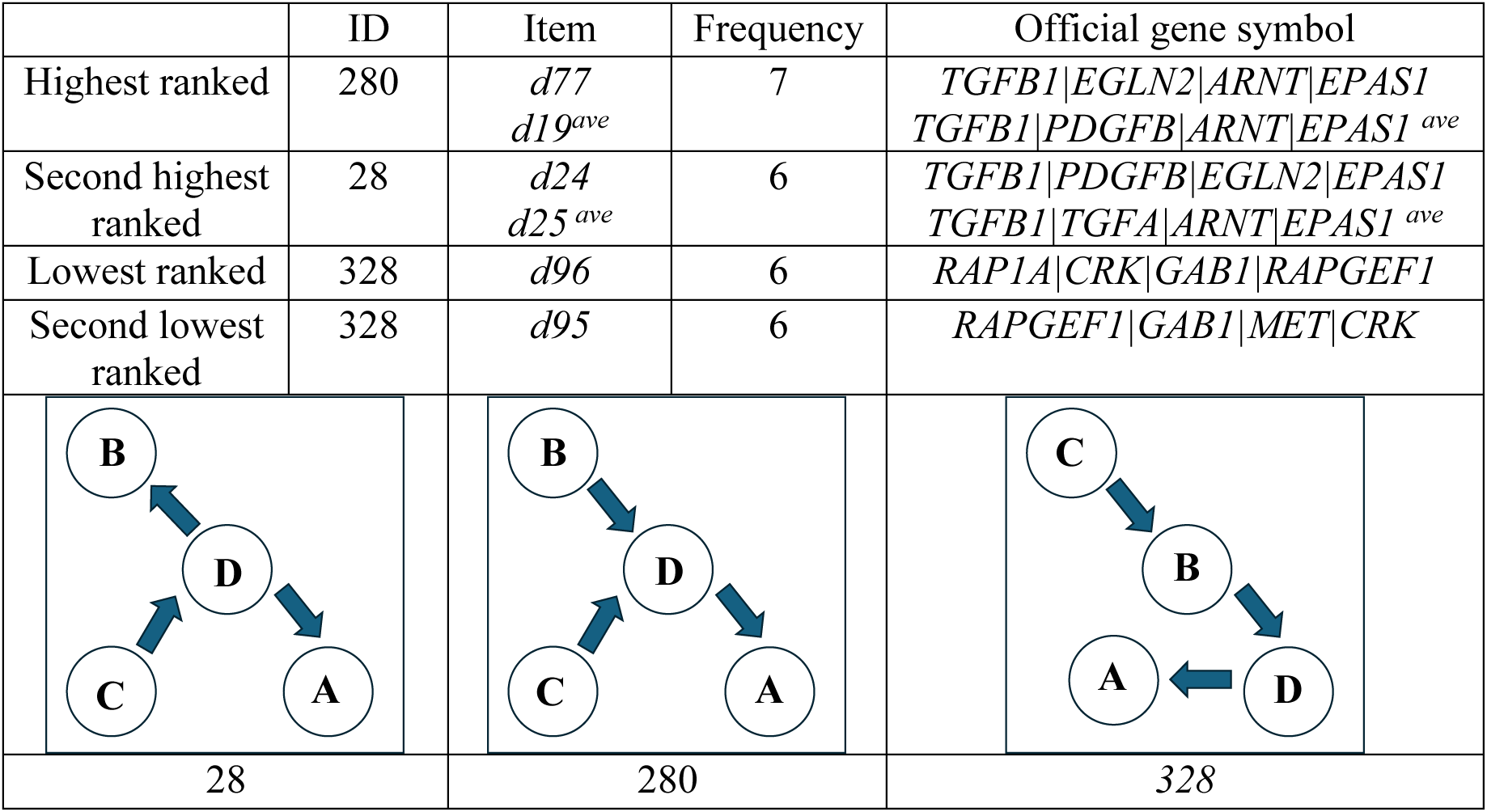
The subgraph IDs and gene symbols of the top two (highest and second highest) and bottom two (lowest and second lowest) ranked 4-node GRMs (both weighted and average *DS* are used for *SC* ranking). The topology of the top two and bottom two ranked 4-node GRMs are shown at the bottom.

When using four features (*FC_HR_Occ_cdg*), both weighted *SC* and average *SC* are applied. For the other 10 combinations, we recalculated the weighted score values for each combination using the same *DS* values derived from the *FC_HR_Occ_cdg* combination for ranking (Supplementary Table 2). The results of *the highest and lowest five ranked GRMs for the 11 combinations* is summarized in Supplementary Table 3.

An alternative method for selecting the top and bottom ranked GRMs is to identify GRMs that most frequently appear as the highest-ranked and second highest-ranked across the 11 combinations. The results show that item *d24* appears most frequently as the highest-ranked (frequency = 3), and item *d21* as the second-highest-ranked (frequency = 2). Notably, the above-mentioned items differ from the items *d77* and *d18* listed in Table 2. Among the lowest-ranked items across 11 combinations, *d95* appears three times, while *d99* and *d108* each appear twice. Notably, *d95* consistently ranks lowest in this context.

### To assess the relevance of the GRMs related to renal cancer using Hallmarks of cancer

Supplementary Table 4 summarizes the top three significant cancer hallmark annotations (p-value < 0.05) for the top two and bottom two 4-node GRMs. While the p-values for these cancer hallmark annotations exceed 0.05 in some cases, a clear distinction between the top two and bottom two GRMs emerges.

The top two GRMs show a direct association with cancer pathways, while only the second-lowest ranked GRM displays a significant association with cancer hallmark formation. The lowest-ranked GRM, however, lacks any significant hallmark annotation, further reinforcing that our method is capable of distinguishing GRMs related to tumor metastasis from other network components within the KIRC network.

When applying an alternative method to select the top (*d24, d21*) and bottom (*d95, d99, d108*) ranked GRMs, the hallmark annotations also highlight a notable difference between the highest two ranked and lowest two ranked GRMs. Supplementary Tables 4 and 5 present nearly identical findings, with the addition of “IL6 JAK STAT3 SIGNALING” and “COAGULATION” found in Supplementary Table 4. This consistency between both scoring methods confirm the robustness and reliability of the results.

### Further assessment of the GRMs’ relevance to KIRC using GOEA

Supplementary Table 6 provides a summary of the highest two and lowest two ranked 4-node GRMs, including their enriched pathways (Wikipathwayss annotations) and associated diseases (DisGeNET annotations). These annotations reflect a significant biological difference between the top two and bottom two ranked GRMs. Specifically, the top two GRMs are closely related to renal cell carcinoma and cancer pathways, while the bottom two modules predominantly show annotations that are unrelated to cancer formation, with only the second annotation being associated ‘MET in type 1 papillary renal cell carcinoma’. Supplementary Files 3 and 4 provides a comprehensive list of the results of GOEA for the top two and bottom two ranked GRMs.

Among the DisGeNET annotations, the highest-ranked GRM reveals a connection to hereditary paraganglioma-pheochromocytoma (PGL/PCC) syndromes, which involve tumors that grow along the spine or adrenal glands. PGL/PCC is characterized by paragangliomas, often found at the carotid artery split in the upper neck. In contrast, the second-highest ranked GRM is linked to glioblastoma multiforme and hereditary PGL/PCC syndrome, highlighting a strong association with tumorigenesis.

The bottom two ranked GRMs, however, exhibit weaker tumor-related annotations. While the lowest ranked GRM shows no connection to tumors, the second-lowest GRM reveals associations with neoplasm invasiveness (the ability of tumors to invade surrounding tissue) and squamous cell carcinoma (skin cancer).

This suggests that the top-ranked GRMs are significantly more associated with tumor formation than the bottom-ranked GRMs, further validating the effectiveness of our method in isolating tumor metastasis-associated GRMs.

We also calculated *JI* to compare the top and bottom ranked GRMs for the following two scenarios (i) we re-calculate the *CD* and *DS,* (ii) we use the original *CD* and *DS*. This was done to investigate whether the ranking of GRMs is robust across variations in the analysis method. The *JI* values for the top five and bottom five ranked GRMs were computed for both “different *CD, DS*” and “same *CD, DS*” scenarios. Interestingly, the *JI* values indicated minimal change in the GRM rankings when moving from five to ten items, suggesting that the ranking method is stable and consistent.

*JI* calculation for the top 5, bottom 5 data items for (i) different *CD*, *DS,* (ii) same *CD, DS*. Repeat that *JI* for top 10. bottom 10. Then we can see if the *JI* changes a lot from 5 to 10 items, if there is not much change this suggests that the use of (i) and (ii) does not affect the ranking or its robustness.

To further validate the CFA method, we considered GRMs composed of three genetic elements (3-node GRMs). The results of the GOEA also showed a disparity between the top-ranked GRMs and the bottom-ranked GRMs (Supplementary Table 7, Supplementary File 6). Again, the top two GRMs are closely related to renal cell carcinoma and cancer pathways, while the bottom two GRMs predominantly show annotations that are unrelated to cancer formation, with only the lowest ranked GRM associated ‘MET in type 1 papillary renal cell carcinoma’. This reinforces the validity of the CFA method, as the results align with those derived from 4-node GRMs.

### Relevance of GRMs to drug targets of KIRC cohort

In Table 3, we list drugs associated with the treatment of renal cancer or anemia linked to chronic renal disease. A striking finding is that four out of five genes (80%) targeted for renal cancer treatment are linked to the top two ranked GRMs, reinforcing the clinical relevance of our identified modules. Furthermore, we found no drug associations for the bottom two ranked GRMs, further indicating that the top GRMs have stronger potential for therapeutic applications.

**Table 3.**
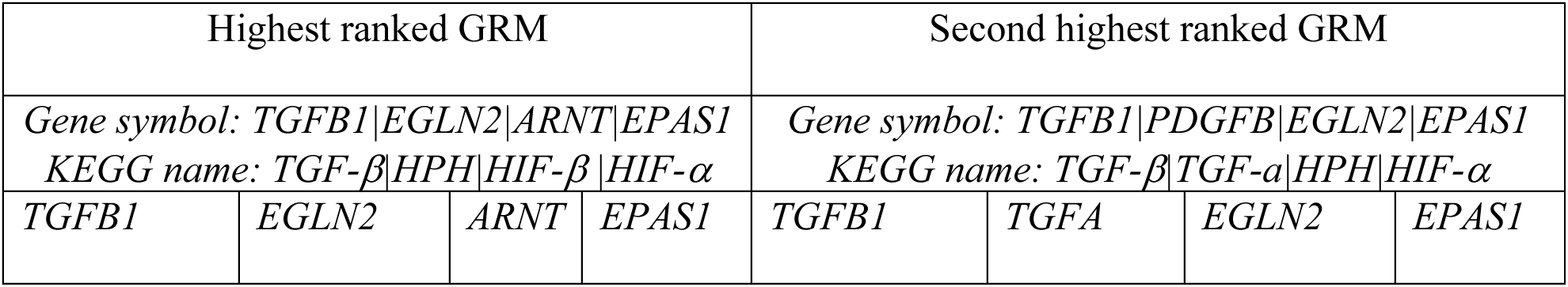
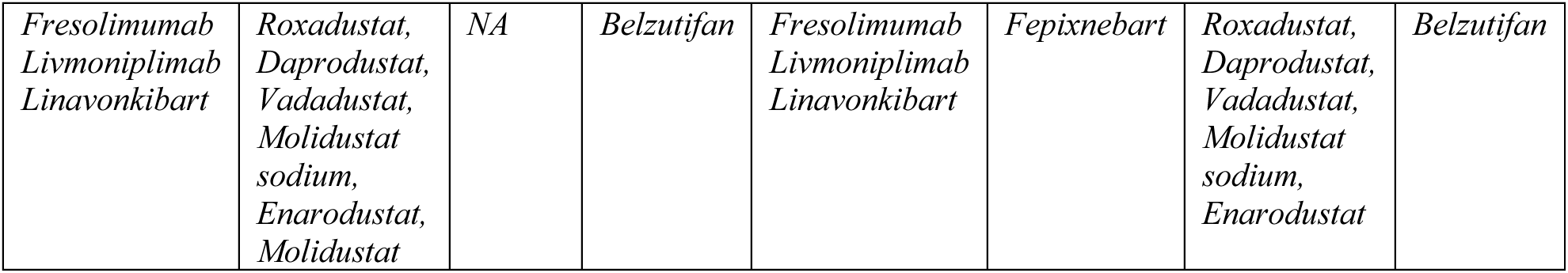
The drugs (USAN/INN/TN) recorded by the KEGG drug database that are used for the treatment of renal cancer or anemia associated with chronic renal disease.

### Survival analysis of KIRC cohort using the UALCAN database

We conducted survival analyses to study the gene cooperativity of the identified GRMs and compared them with existing findings. The first analysis was performed using the UALCAN database, and the second utilized the cBioPortal database [61]. The results of the survival analysis using the UALCAN database [59, 60] are shown in Figure 3, highlighted that high expression of *TGFB1* and low expression of *ARNT, EPAS1, TGFA*, and *PDGFB* are significantly associated with lower survival rates in KIRC patients. Specifically, five out of six genes (83.3%) identified as risk biomarkers show a negative impact on survival, while *EGLN2* (Figure 3 (F)) did not show any significant survival impact. Conversely, among the eight genes in the five lowest-ranked GRMs, only one gene, *SOS1*, impacted survival probability (Supplementary Figure 1). These findings further emphasize that the top-ranked GRMs are more closely associated with survival outcomes and tumor metastasis.

**Figure 3.**
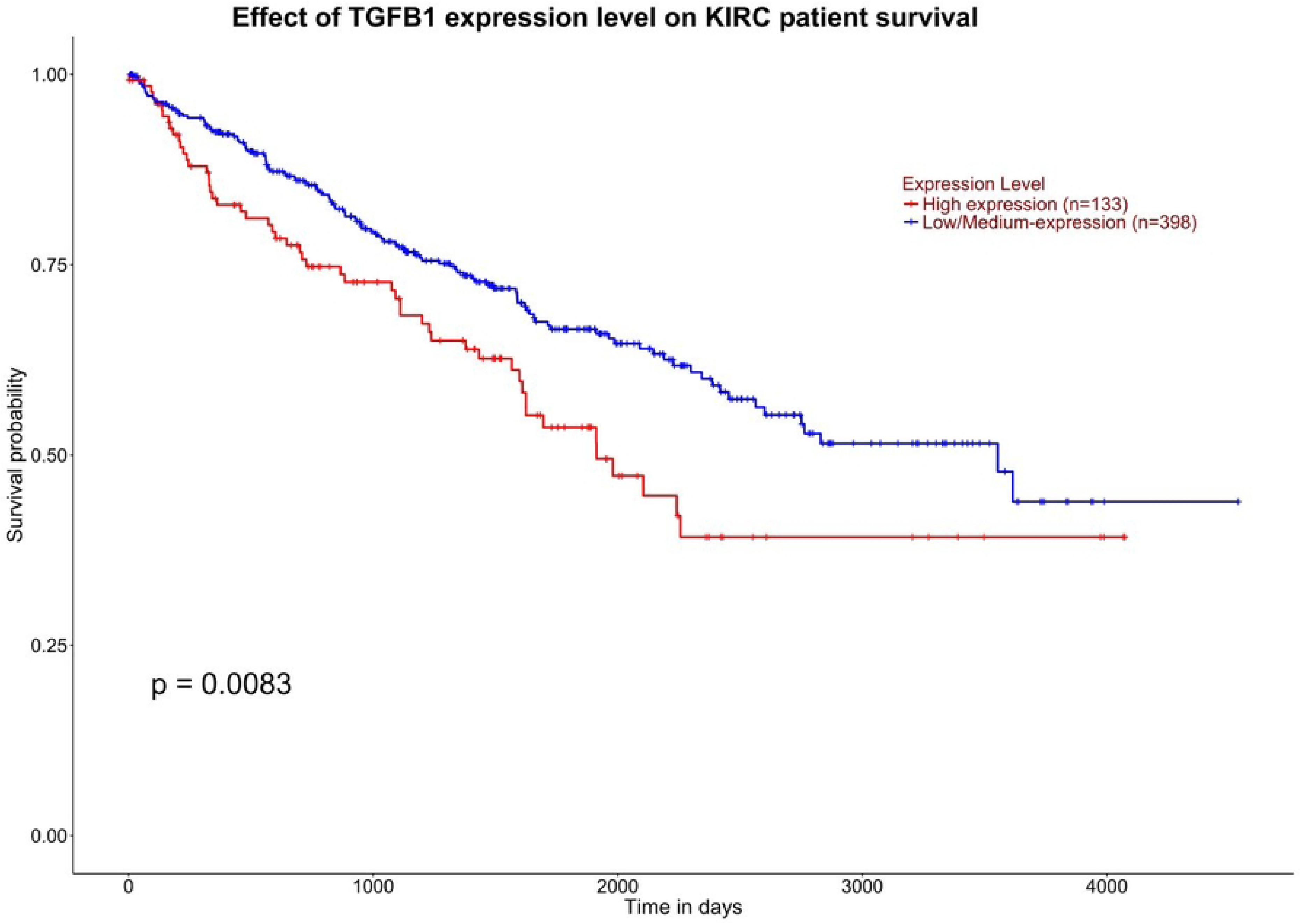
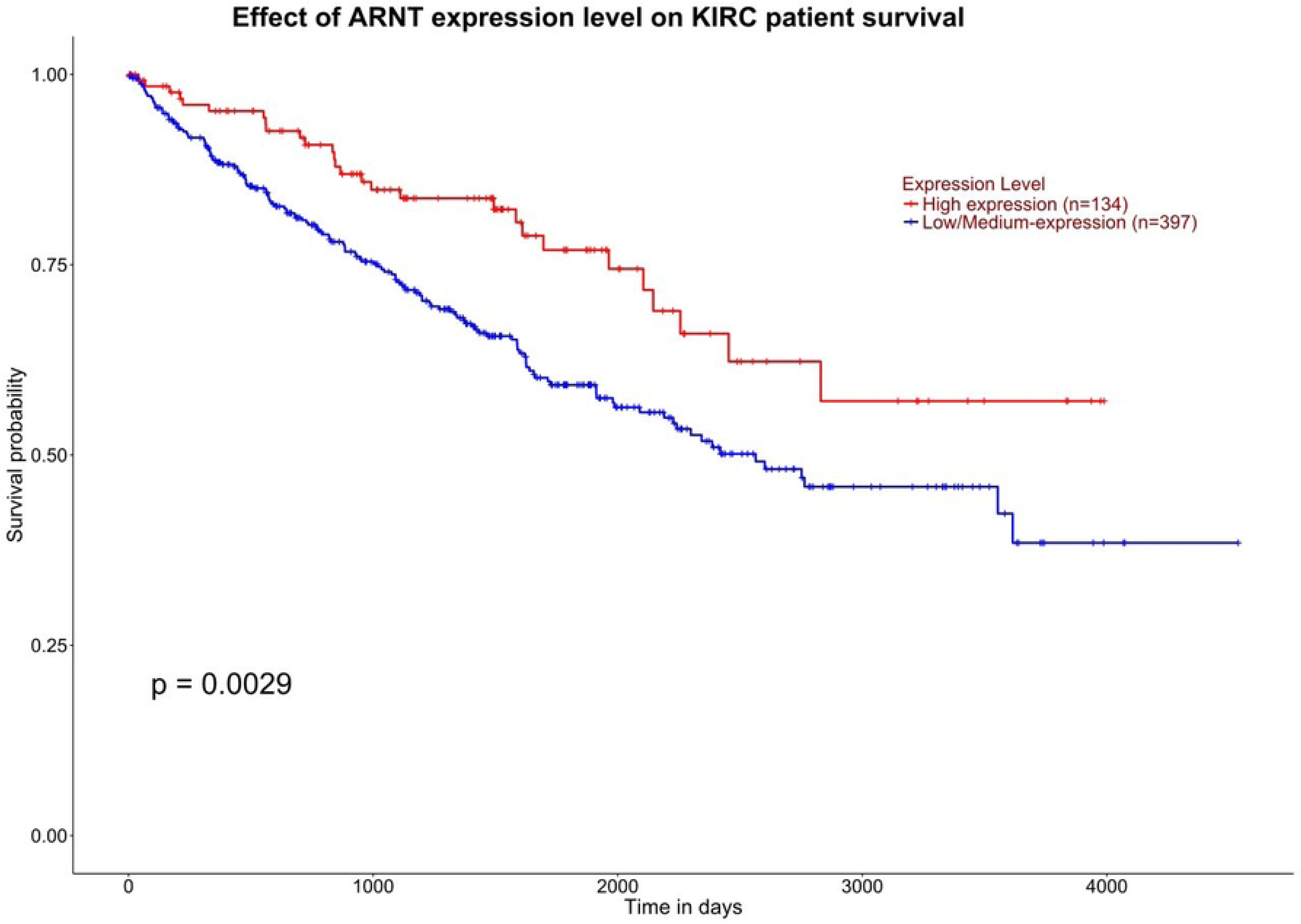
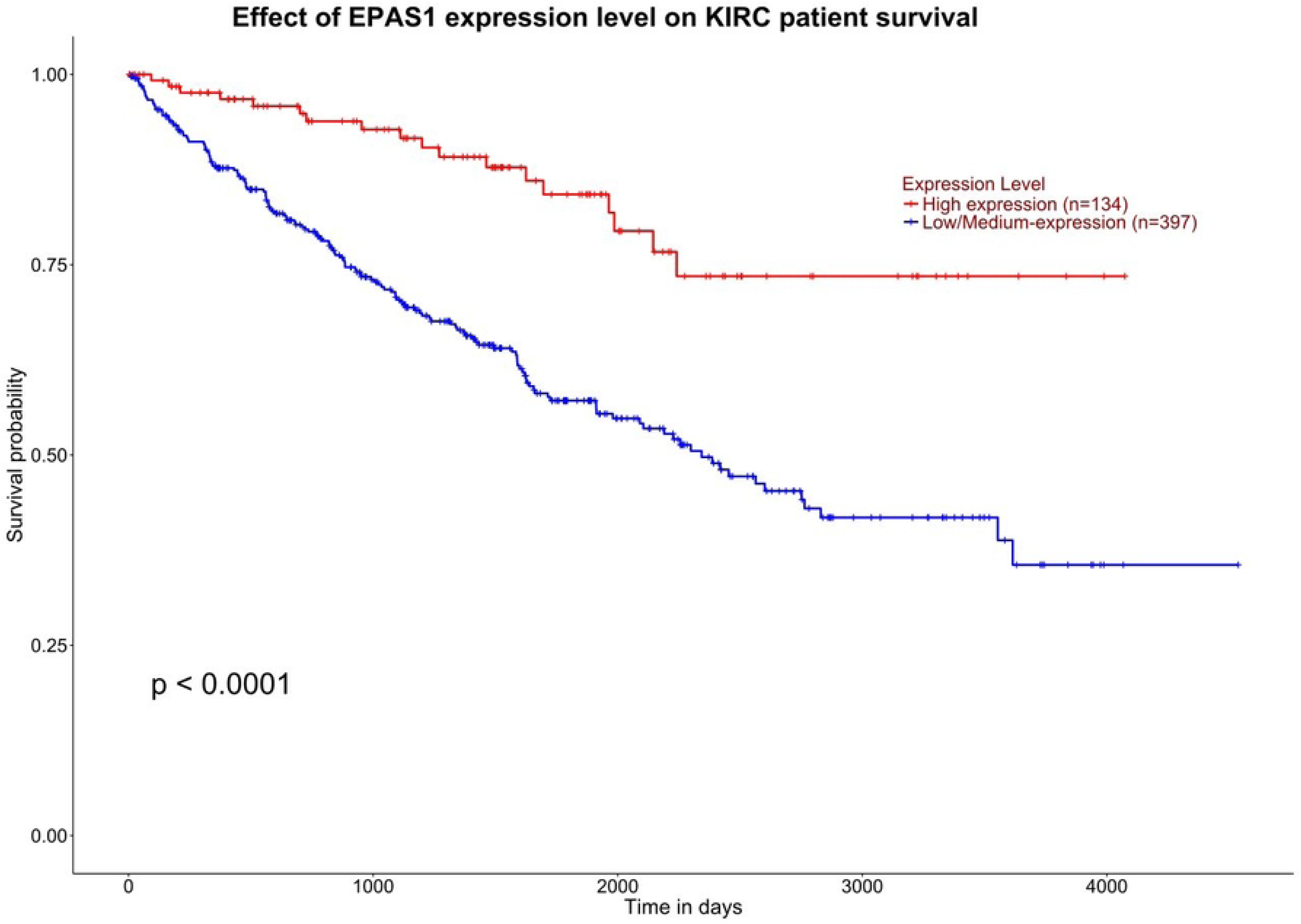
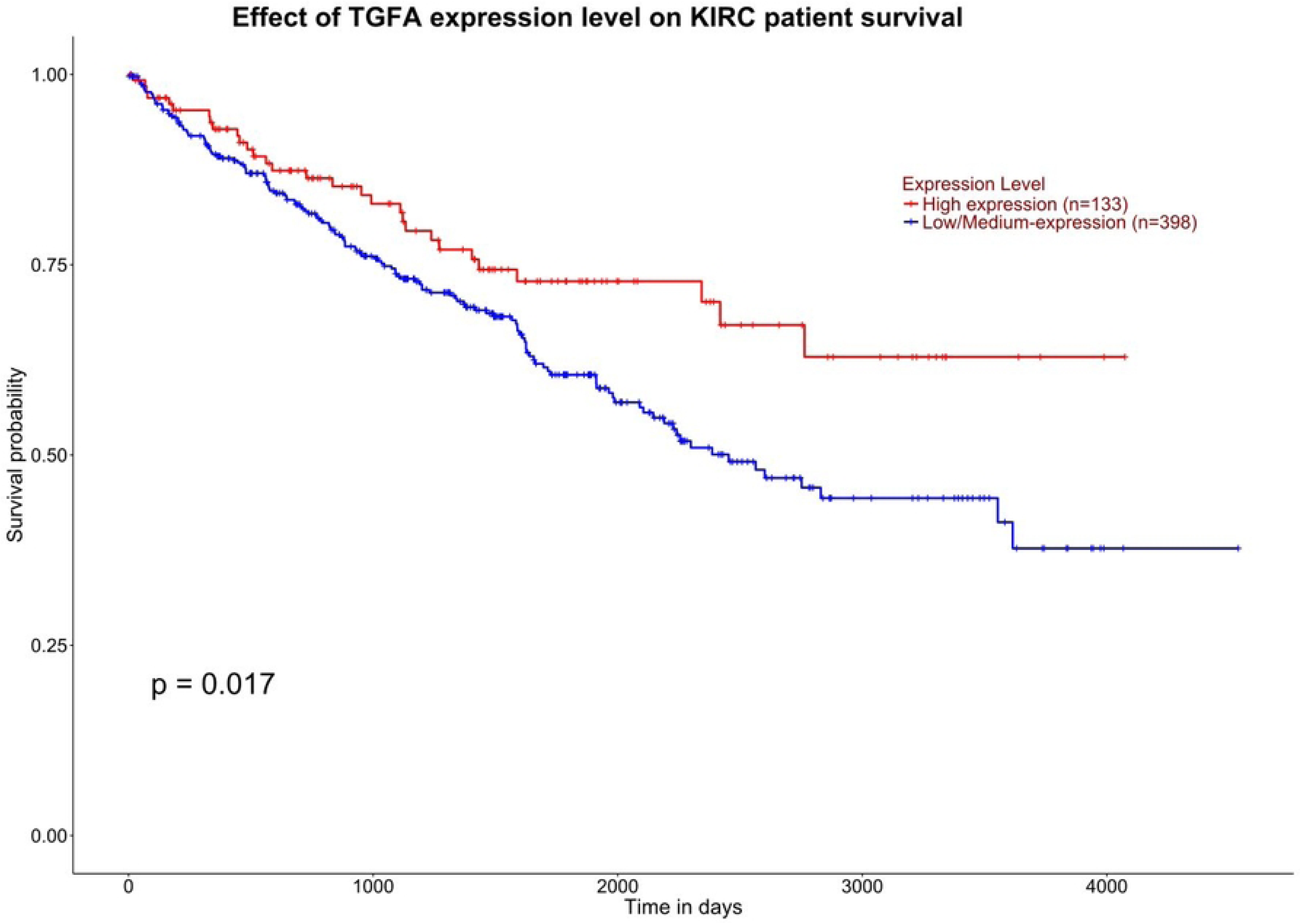
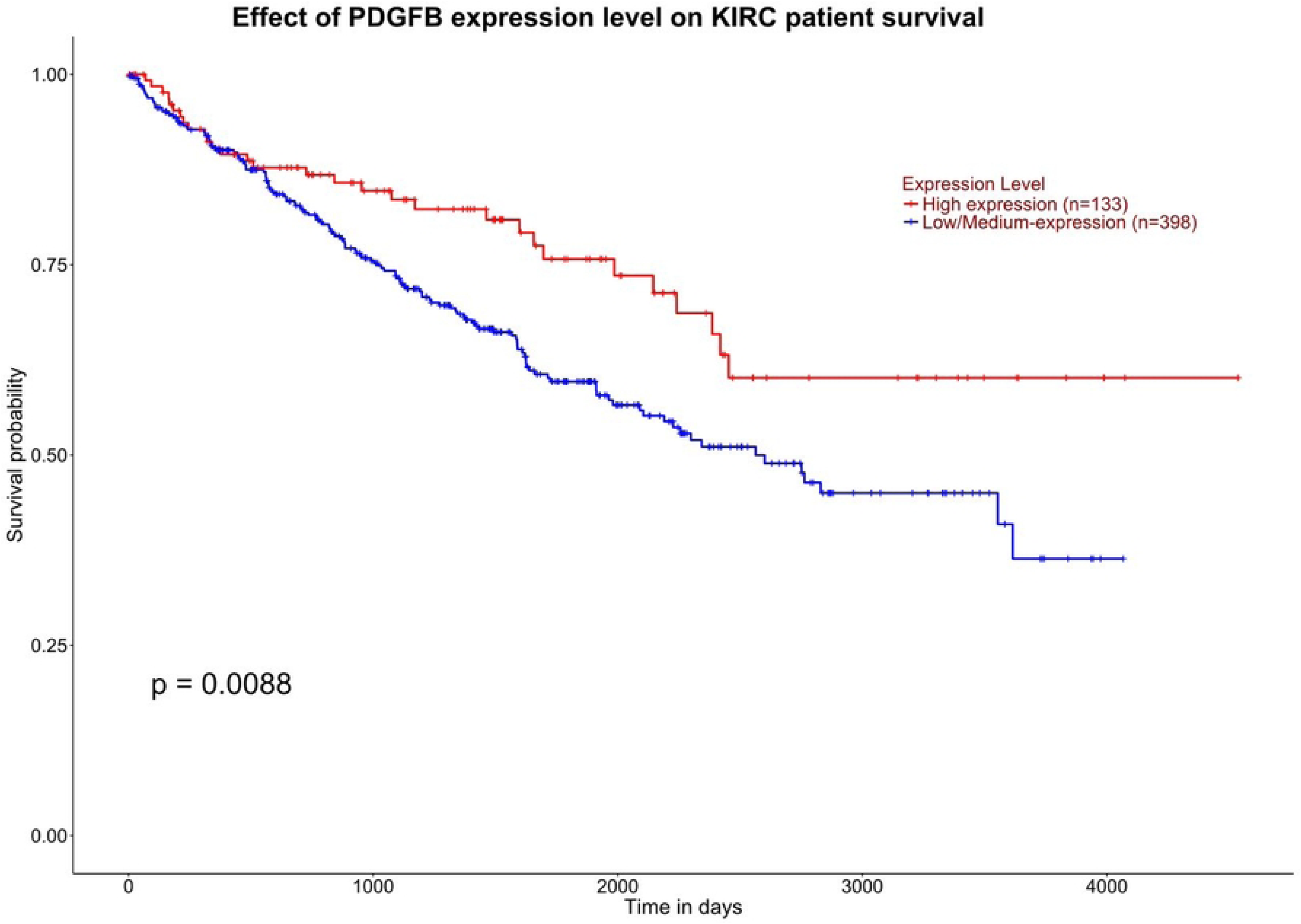
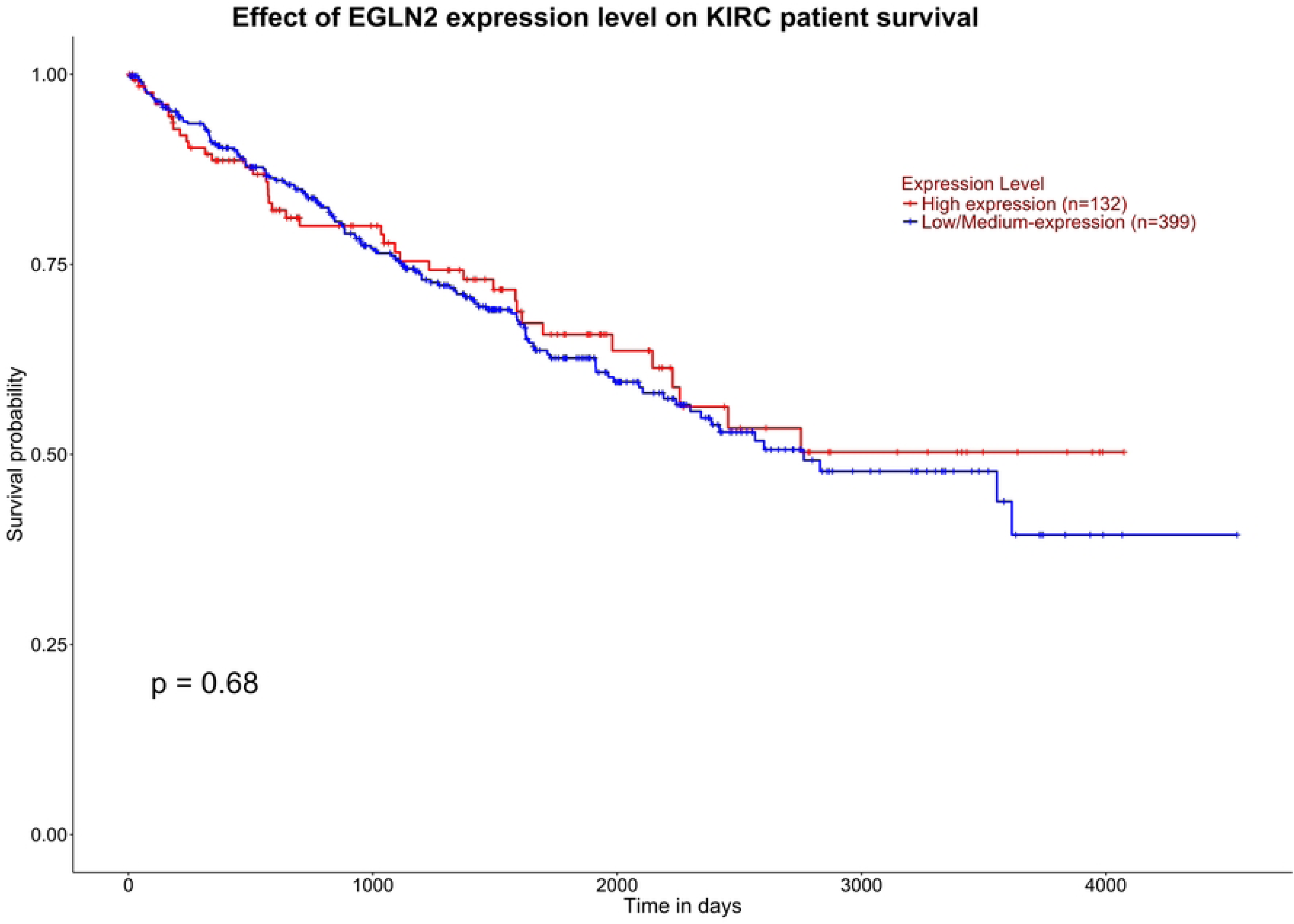
The results of the survival probability analyses, (A) *TGFB1* (B) *ARNT, (C) EPAS1* (D) *TGFA, (E) PDGFB* and (F) *EGLN2.* There is statistically significant negative impact on survival probability, leading to lower survival rates.

### Survival analysis of KIRC cohort using the cBioPortal databases and metastasis-associated genes (*Occ*)

The results of the survival analysis using the highest two and lowest two GRMs, conducted through the cBioPortal database, are shown in Table 4. We tested the consistency of the results by repeating the analysis three times with different TCGA datasets: (i) TCGA Firehose Legacy, (ii) TCGA Nature 2013, and (iii) TCGA GDC.

**Table 4.** Survival analysis was conducted using the cBioPortal database for the two highest-and two lowest-ranked GRMs, as well as four additional genes from reference [18]. The symbols ‘Y’ and ‘NS’ indicate p-values of <0.05 and >0.05, respectively. Genes related to metastasis are denoted by the abbreviation ‘*Occ’*.

| Genes | Official gene symbol | Firehose | Nature2013 | GDC | Occ |
| --- | --- | --- | --- | --- | --- |
| Highest ranked | *TGFB1, PDGFB, ARNT, EPAS1<sup>ave</sup>* | Y | Y | Y | 11 |
| Second highest ranked | *TGFB1, TGFA, ARNT, EPAS1<sup>ave</sup>* | Y | Y | Y | 11 |
| Lowest ranked | *RAP1A, CRK, GAB1, RAPGEF1* | Y | Y | NS | 9 |
| Second lowest ranked | *RAPGEF1, GAB1, MET, CRK* | Y | Y | NS | 8 |
| metastasis-associated genes [18] | *COL1A1, DCN, FBLN1, POSTN* | NS | NS | NS | 7 |
|  | *COL1A2, DCN, FBLN1, POSTN* | NS | NS | NS | 6 |
|  | *COL5A1, DCN, FBLN1, POSTN* | Y | NS | NS | 4 |
|  | *COL6A3, DCN, FBLN1, POSTN* | Y | NS | NS | 4 |
|  | *COL1A1, COL5A1, COL6A3, DCN* | Y | NS | Y | 4 |
|  | *COL1A1, COL5A1, COL6A3, FBLN1* | NS | Y | NS | 4 |
|  | *COL1A1, COL5A1, COL6A3, POSTN* | Y | NS | Y | 5 |
|  | *COL1A1, COL1A2, COL5A1, COL6A3* | Y | NS | Y | 5 |
| metastasis-associated genes [62] | *VHL, PBRM1, BAP1, SETD2* | NS | NS | NS | 7 |
| metastasis-associated genes [63] | *ABCG1, HAVCR2, CD14, TGFA* | NS | S | NS | 7 |

The Genomic Data Commons (GDC) utilizes harmonized clinical data, which may alter survival times, censoring status, or event definitions. The other two datasets might have outdated or less complete clinical follow-up, which can inflate or deflate significance.

We emphasize that our survival analysis was performed using a combination of four biomarkers, rather than a single one. This analysis fundamentally differs from methods that rely on four features— *logFC*, *HR, Occ* and *cdg* —which are attributes of individual biomarkers. Thus, our study highlights the cooperative effect of a set of biomarkers, which should not be equated with the effects of individual biomarkers.

Additionally, we compared our findings with 3 independent KIRC cohort studies, which identified seven genes [18]— namely, the collagen family (*COL1A1, COL1A2, COL5A1*, and *COL6A3*), *DCN, FBLN1, and POSTN*—that were significantly upregulated in metastatic tumors compared to primary tumors. Another study [62] reported four genes— *VHL*, *PBRM1*, *BAP1*, *SETD2* —as key drivers of tumor initiation and metastasis. The third study by [63] identified increased mRNA expression of four genes— *ABCG1, HAVCR2, CD14, TGFA*— in KIRC samples for prognostic genes, relative to adjacent control tissue.

Table 4 summarizes the survival analysis results using the cBioPortal database for the top two and bottom two ranked GRMs (derived from average *SC*, as shown in Table 2), along with four genes selected from the seven genes [18], which include one gene from the collagen family and *DCN, FBLN1,* and *POSTN*.

Different combinations of the seven genes were tested to identify which sets of four genes yield consistent survival analysis results. First, one gene from the collagen family was selected and combined with the three non-collagen genes (*DCN, FBLN1,* and *POSTN*). Second, a different collagen gene was chosen and again combined with the same three non-collagen genes. Third, three collagen genes (*COL1A1, COL5A1, COL6A3*) were each combined with one of the non-collagen genes to form three additional gene sets.

Table 4 presents the survival analysis results. The top two ranked GRMs, based on average SC, consistently produced statistically significant p-values (<0.05) across all three TCGA datasets. In contrast, the bottom two ranked GRMs had at least one non-significant (NS) result. Gene sets selected from Reference [49] also showed mostly NS results, with at least two NS outcomes observed in four out of eight cases. It was also found that the key drivers of tumor initiation and metastasis yield NS results in survival analyses across all three TCGA datasets. Furthermore, genes related to metastasis, ‘*Occ’*, are relative fewer compared to the highest two ranked GRMs. We noted that the lowest two ranked GRMs have a moderate number of ‘*Occ’*, this could explain why the survival analysis difference between the highest two and lowest two are moderate.

We compared our findings with three independent KIRC cohort studies. Survival analysis, conducted using a combination of four biomarkers, indicates that our current approach is more effective in identifying GRMs associated with metastasis.

Ref 56 → macrophage differentiation–associated genes (MDGs)

### Quantifying the difference for the weighted *SC* and average *SC* results using Jaccard index

To quantify the differences in GRM rankings derived from using weighted *SC* and average *<u>SC</u>* methods, we calculated the *JI* for various subsets of the ranked GRMs. For the top five, bottom five, top ten and bottom ten ranked GRMs, we observed that both methods identified a similar set of GRMs, with *JI* values of 2/3 for all these subsets, except *JI* (bottom 10) = 9/11. This suggests that the weighted *SC* and average *SC* methods produce nearly identical results for these GRM rankings.

The *JI* analysis for additional combinations (provided in Supplementary File 5) showed that for most cases, the *JI* was exactly one for the top five and bottom five ranked GRMs, indicating complete agreement between the weighted *SC* and average *SC* methods. This further underscores the reliability of the results obtained using both methods.

### To assess the relevance of the 3-node GRMs related to renal cancer

In addition to the 4-node GRMs, we also examined 3-node GRMs (Supplementary Table 7) to assess whether the top and bottom ranked GRMs demonstrated distinct biological functions. The comparison between the 3-node and 4-node GRMs (Supplementary Table 7 and Table 2) revealed that the 4-node GRMs encompass all the 3-node GRMs, except for the second-lowest ranked 4-node GRM, where the gene *RAP1A* is absent. This suggests a high level of consistency between the results of the 4-node and 3-node GRMs, further supporting the validity of our method in identifying metastasis-associated GRMs.

## Conclusion

We have tested our hypothesis, which suggests that the 4-node GRMs consisting of genetic elements are key candidates for metastasis-associated gene modules. By developing an integrated approach that combines the subgraph method and CFA, we uncovered metastasis-associated GRMs. Validation through enrichment analysis, drug–target gene insights, survival data, and comparison with previously published work demonstrates its potential for identifying metastasis-associated target genes and discovering therapeutic drug candidates.

### Limitation and future works

Despite the promising results, some limitations must be acknowledged. One key limitation is that our approach does not consider the rank combination method due to the constraints imposed by the cancer driver genes (*cdg*) scoring method, which allows only a limited number of ranks. To address this issue, we suggest incorporating additional genetic categories, such as passenger driver genes and essential genes, into the scoring system to refine the evaluation. Additionally, expanding the number of metastasis databases beyond three could help mitigate the issue of having too many tiers.

Furthermore, incorporating external datasets, such as Firebrowse (http://firebrowse.org/), could further improve the ranking accuracy. Firebrowse offers access to comprehensive TCGA data, including detailed cancer staging information, which could enrich our analyses and provide more robust results.

Future research could also explore the scalability issue in constructing larger GRMs. For instance, multiple 3-node GRMs could be merged to construct n-node GRM, addressing scalability challenges and identifying larger metastasis-associated GRMs.

Finally, the findings suggest that the developed method could be applied to predict the relevance of new GRMs inferred from the KIRC cancer cohort. By calculating ranks using the four genetic component scores—*FC, HR, cdg*, and *Occ*—GRMs can be ranked and assessed for their potential association with tumor metastasis, providing valuable insights for future cancer research and drug development.

Our comprehensive analysis demonstrates that the method developed for ranking and evaluating GRMs in the context of KIRC is effective in isolating tumor metastasis-associated GRMs. By integrating multiple scoring methods and using GOEA, hallmark annotations, drug-target information, and survival data, we provide a robust framework for identifying significant GRMs related to cancer formation and metastasis.

**Supplementary information –** The file consists of

i. Supplementary Figure 1 - The results of the survival probability analyses,
ii. Supplementary Table 1 - The results of the *DS* score for the 11 possible scoring method combinations,
iii. Supplementary Table 2 - A comparison of the results using weighted and average *DS* for ranking the GRMs,
iv. Supplementary Table 3 - The results of the highest and lowest five ranked GRMs for the 11 combinations,
v. Supplementary Table 4 - The top three most significant cancer hallmark annotations based on using weighted score combination and average score combination,
vi. Supplementary Table 5 - The top three most significant cancer hallmark annotations based on using based on their frequency of being ranked first or second across the 11 combinations.
vii. Supplementary Table 6 - The results of the first two most significant Wikipathways and DisGeNET annotations for the top two and bottom two ranked <u>4-node</u> GRMs,
viii. Supplementary Table 7 - The results of the first two most significant Wikipathways and DisGeNET annotations for the top two and bottom two ranked <u>3-node</u> GRMs.

**Supplementary File 1** - The set of 4-node GRMs identified in the renal kidney cancer network.

**Supplementary File 2** - The complete set of all 199 possible 4-node gene regulatory modules.

**Supplementary File 3** – The results of enrichment analysis for the highest two ranked 4-node GRMs across all 11 combinations of the four scoring methods.

**Supplementary File 4** – The results of enrichment analysis for the lowest two ranked 4-node GRMs across all 11 combinations of the four scoring methods.

**Supplementary File 5** – Results of the Jaccard index (*JI*) between the weighted and average score approaches, comparing (i) the top five and bottom five GRMs, and (ii) the top ten and bottom ten GRMs, across all 11 combinations of the four scoring methods.

**Supplementary File 6** – The results of enrichment analysis for the highest two and lowest two ranked 3-node GRMs using four scoring methods.

## Acknowledgments and Funding

Miss Aninda Astuti and Dr. Ka-Lok Ng works are supported by (i) National Science and Technology Council (NSTC), Taiwan (grant numbers: NSTC 112-2221-E-468-021, 113-2221-E-468-014, 114-2221-E-468-009), and (ii) Asia University and China Medical University Hospital (grant number: ASIA-112-CMUH-6, ASIA-114-CMUH-07). The funding bodies were not involved in the design of the study, in the collection, analysis of the data, or in writing of the manuscript. URL address of [1] the NSTC https://www.nstc.gov.tw/?l=en, [2] Asia University https://web.asia.edu.tw/ and [3] China Medical University Hospital https://www.cmuh.cmu.edu.tw/Home/CmuhIndex_EN?lang=1.

We would also like to thank Dr. Efendi Zaenudin for preparing Supplementary File 2, which contains the 199 patterns of the 4-node GRMs

## Author contributions

Aninda Astuti is responsible for various duties, including data-related programming, conducting formal analyses, prepared figures, and validation. Christina Schweikert and D. Frank Hsu offer expertise in formal CFA analysis and contribute to the manuscript review. Ka-Lok Ng conceptualized the research, interpreted the results, edited the manuscript, and obtained financial support.

## Data availability statement

The raw data is derived from analyzing the publicly available KEGG database, resulting in the four-node GRM module, which is provided in Supplementary File 1. The code used for analyzing the GRM is publicly available on GitHub at the following link: https://github.com/anindaastuti/GRMs_CFA

## Declaration of generative AI and AI-assisted technologies in the writing process

During the preparation of this work, the author, Ka-Lok Ng, used ChatGPT and Copilot for grammar correction, and content revision of our English writing. After using this tool/service, the author, Ka-Lok Ng, reviewed and edited the content as needed and take full responsibility for the content of the publication.

